# Reachability-Preserving Minimum Edge Cut Problem and Applications in Biology

**DOI:** 10.64898/2026.06.01.729192

**Authors:** Jing Xie, Qi Duan

## Abstract

Biological pathway analysis often requires identifying interventions that block reachability to an undesirable state, such as a disease-associated module, toxic byproduct, or adverse phenotype, while preserving reachability among essential biological functions. Motivated by this setting, we study the Reachability Preserving Minimum Edge Cut (RPMEC) problem: given protected terminals *s*_1_ and *s*_2_ and a target terminal *t*, the goal is to remove a minimum-cost set of edges that separates *s*_1_ and *s*_2_ from *t* while keeping *s*_1_ and *s*_2_ connected. This formulation naturally models pathway-level intervention design, where one seeks to disrupt harmful signaling, metabolic, or interaction routes without breaking required functional connectivity. We revisit the three-terminal undirected edge-cut case and analyze a Dijkstra-style dynamic programming algorithm that is exact on planar graphs but fails on general graphs. We characterize the structural condition required for exactness, namely frontier-realizability of optimal source-side regions, and identify biological graph representations where this condition is likely to hold after appropriate preprocessing, including curated planar pathway maps, Reactome-style hierarchy trees, SCC-contracted feedback modules, metabolic building-block DAGs with dominator structure, and functional-module quotients of protein interaction networks. We further discuss directed variants, approximation strategies, and exact alternatives based on ASP, MILP, bounded-treewidth dynamic programming, and important separators. The results provide a graph-theoretic foundation for deciding when fast greedy computation is reliable for biological pathway intervention problems and when more expressive exact optimization methods are needed.

**Author Summary:** Many real-world networks require interventions that separate harmful or undesirable states while preserving essential connectivity. This situation appears in biological pathway analysis, where one may want to block reachability to a disease-related module, toxic byproduct, or adverse phenotype without disrupting communication among essential genes, proteins, reactions, or metabolites. We study this problem through the Reachability Preserving Minimum Edge Cut formulation. Unlike ordinary minimum cut, the solution must satisfy both a separation requirement and a preservation requirement. We show why a natural Dijkstra-style algorithm works only under specific structural conditions, such as planar, laminar, or module-like pathway graphs, and why it may fail on general graphs. The results help identify when fast graph algorithms are reliable for biological intervention problems and when exact optimization tools such as Answer Set Programming or integer programming are more appropriate.

## Introduction

Minimum cut is one of the central problems in graph algorithms. Given two terminals, a classical cut problem asks for a minimum-cost set of edges whose deletion separates them, and the standard max-flow/min-cut framework gives efficient polynomial-time algorithms for this task [1]. Many modern applications, however, require more than separation. In a network-defense instance, one may want to isolate an attacker-controlled node while keeping critical services mutually reachable. In a biological pathway instance, one may want to block reachability to a disease-related or toxic state while preserving reachability among essential genes, proteins, reactions, or metabolites. These examples motivate a cut problem with both a negative requirement, namely separating a bad destination, and a positive requirement, namely preserving important reachability on the source-side.

This paper studies such a problem, which we call the *Reachability Preserving Minimum Edge Cut* (RPMEC) problem. In the basic undirected three-terminal form, the input is a connected weighted graph *G* = (*V, E*) with terminals *s*_1_, *s*_2_, and *t*. The goal is to remove a minimum-cost edge set *F* ⊆ *E* such that *s*_1_ and *s*_2_ remain connected in *G* − *F*, while *t* is disconnected from their component. Equivalently, one may choose a connected source-side vertex set *X* with

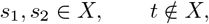

and minimize the weight of the boundary *δ*(*X*). This is the edge-cut version of the connectivity-preserving minimum cut problem studied in [2], and it is also a special case of the broader cut-uncut family, where some terminal pairs must be separated while other terminal pairs must remain connected [3, 4]. We use the name RPMEC to emphasize that the essential constraint is preservation of reachability among protected terminals.

A natural approach to RPMEC is the Dijkstra-style dynamic programming algorithm proposed for the three-terminal planar case in [2]. Starting at *s*_1_, the algorithm repeatedly computes minimum cuts that force a previously certified source-side component together with one new frontier vertex, and then settles the cheapest such component. This resembles Dijkstra’s shortest-path algorithm: a growing settled region is expanded by the cheapest local relaxation. The analogy is powerful, but it is also delicate. In shortest paths, a globally shortest unsettled path always has a predecessor that has already been settled. In RPMEC, the analogous statement is that the next optimum source-side cut region must be visible through a one-frontier cut relaxation. This visibility property holds in planar instances by noncrossing structure, but it need not hold in general graphs.

Figure 1 gives a smaller counterexample with small integer weights. The source-side set *{s*_1_*}* has cut value

**Fig 1.**
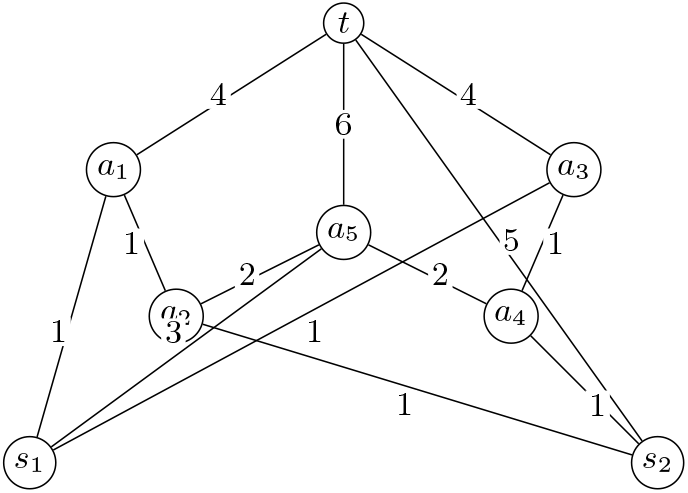
A smaller counterexample for the undirected Dijkstra-style RPMEC algorithm. The algorithm settles the left/right branches first and returns a cut of value 16, while the true optimum has value 15.

**Fig 2.**
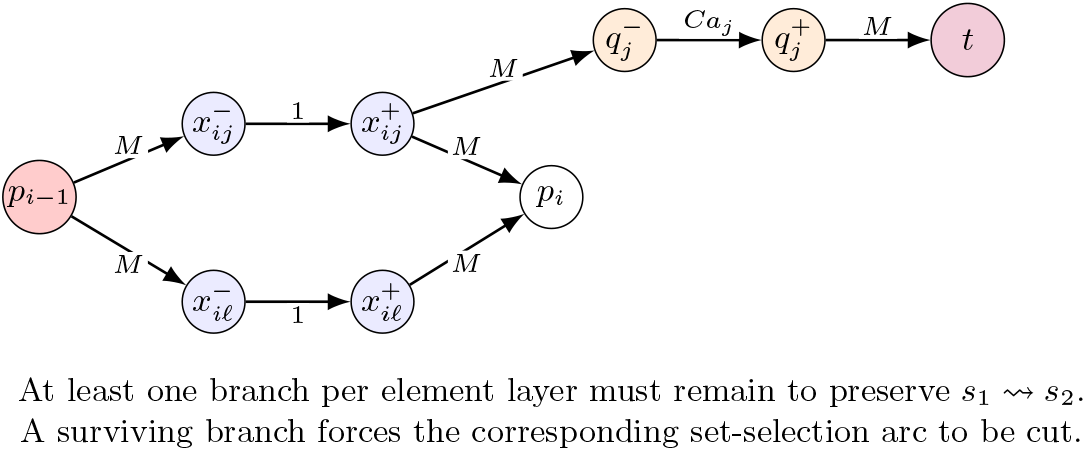
One layer of the DAG reduction from Set Cover. Each surviving branch represents choosing a set that covers the corresponding element.

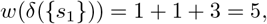

so initially the algorithm starts from 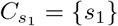.

For the first frontier step, the three possible relaxations are

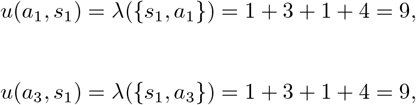

and

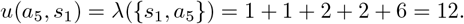

Hence the Dijkstra-style algorithm settles *a*_1_ and *a*_3_ before *a*_5_.

Next, after settling the left and right upper branches, the relaxations to *a*_2_ and *a*_4_ are

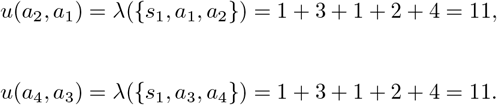

These are still smaller than *u*(*a*_5_, *s*_1_) = 12, so the algorithm settles *a*_2_ and *a*_4_ before it settles *a*_5_.

When the algorithm finally extends from the left side to *s*_2_, it obtains the source-side component

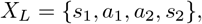

whose cut value is

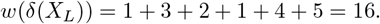

By symmetry, the right branch gives the same value 16.

However, the true optimum is the mixed component

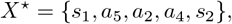

whose cut value is

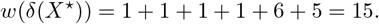

Therefore the Dijkstra-style algorithm returns value 16, while the true RPMEC optimum is 15.

The reason for the failure is that the optimal source-side component *X*^⋆^ uses the middle vertex *a*_5_ together with both lower branch vertices *a*_2_ and *a*_4_. This mixed component is not exposed by any single frontier relaxation from the components settled earlier, so the algorithm commits to a suboptimal branch-based solution.

The main goal of this paper is to identify precisely when this Dijkstra-style algorithm is reliable. Our central answer is a structural one: the algorithm is exact when the optimum source-side regions are *frontier-realizable*. Informally, at every stage of the algorithm, the cheapest true unsettled optimum region must be obtainable by taking a previously settled source-side component and adding one outgoing frontier vertex, without increasing the optimum cut value. This condition is necessary and sufficient for the label-setting behavior of the algorithm under fixed tie-breaking. It explains why the algorithm works on planar graphs, why it also works on laminar or hierarchical instances, and why it fails on graphs with crossing or interlocking optimum regions.

We also study the directed analogue. In a directed graph, the protected requirement is not merely that *s*_1_ and *s*_2_ lie in the same undirected component, but that *s*_1_ can reach *s*_2_ after the cut. A feasible directed source-side set *X* must satisfy

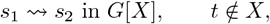

and the cut is the outgoing boundary *δ*^+^(*X*). The directed setting is more subtle because vertex-set overlap does not imply that directed reachability is preserved. Accordingly, the directed Dijkstra-style algorithm requires a stronger condition: optimum regions must be both frontier-realizable and reachability-uncrossable. We show that directed acyclicity alone is not enough; even on directed acyclic graphs, the problem can encode global set-cover-like choices and remains NP-complete.

The structural conditions above are especially useful for deciding when fast algorithms are appropriate on practical graphs. In biological pathway analysis, for example, the Dijkstra-style algorithm is justified for pathway instances whose relevant quotient graph is planar, laminar, arborescent, or dominator-compatible. This includes many curated pathway-map abstractions, Reactome-style hierarchy trees, directed signaling cascades with a unique dominator structure, SCC-contracted metabolic module graphs, and functional-module quotients of protein interaction networks [5–9]. On the other hand, arbitrary protein-protein interaction networks, highly cross-talking signaling networks, and dense metabolic graphs with hub metabolites need not satisfy the frontier-realizability condition. For such instances, an exact declarative model such as Answer Set Programming (ASP) provides a flexible fallback, since ASP can directly encode reachability preservation, separation, and cost minimization [10–12].

The contributions of the paper are as follows.

1. We formalize the RPMEC source-side view for undirected and directed graphs, making explicit the relationship between connectivity preservation, reachability preservation, and ordinary minimum cuts.
2. We characterize the exact condition under which the Dijkstra-style algorithm is label-setting in undirected graphs: every next optimum region must be frontier-realizable.
3. We identify sufficient graph and instance classes for correctness, including planar graphs, laminar optimum-region systems, cut-planar skeletons, and biological module quotients with hierarchical or planar structure.
4. We extend the characterization to directed graphs through directed frontier-realizability and reachability-uncrossing, and we explain why directed acyclic graphs do not by themselves make the problem easy.

## Related Work

### Classical minimum cuts and connectivity-preserving variants

Classical minimum-cut problems ask for a minimum-cost set of edges or vertices separating two terminal sets. The max-flow/min-cut theorem gives a polynomial-time foundation for ordinary two-terminal cuts [1], and the same paradigm underlies a large body of graph-separation algorithms. RPMEC differs from ordinary minimum cut because the cut must satisfy an additional “uncut” constraint: the protected terminals must remain mutually connected or reachable after the removal of the cut edges. This additional constraint destroys the direct max-flow/min-cut reduction and is the source of the algorithmic difficulty studied in this paper.

Duan and Xu [2] consider both node-cut and edge-cut variants of the problem, prove hardness and inapproximability results for the node-cut version, and give a Dijkstra-style dynamic programming algorithm for the edge-cut version on planar graphs. Our RPMEC terminology emphasizes the same preservation requirement from a reachability perspective. The present work revisits the Dijkstra-style algorithm and isolates the structural property that makes it correct: the next optimum source-side region must be visible through a one-frontier relaxation.

RPMEC is also a special case of the broader family of cut-uncut problems. The work in [13] shows the polynomial algorithm for a general case of one versus multiple nodes in planar graphs. In the more general Two-Sets Cut-Uncut problem, one is given two terminal sets *S* and *T*, and the goal is to separate all of *S* from all of *T* while preserving connectivity inside each terminal set. Bentert, et al. gave a randomized fixed-parameter algorithm for Two-Sets Cut-Uncut on planar graphs, with running time exponential in the number of terminals but polynomial in the graph size [3]. A later parameterized-complexity study by Bentert et al. showed that Two-Sets Cut-Uncut remains highly nontrivial even as a feasibility problem and mapped which structural parameters admit FPT, XP, and kernelization results [4]. These works support the central theme of our analysis: adding connectivity-preservation constraints to a cut problem can change the complexity landscape dramatically, even when the ordinary cut problem is easy.

Network Diversion is another closely related cut-uncut problem. It asks for a small edge set whose removal forces all surviving *s*-*t* paths through a specified edge, while preserving at least one such path. Curet introduced the directed version in an operations-research setting [14], and Cullenbine et al. developed theoretical and computational results for directed and planar cases [15]. Recent planar-network-diversion work gives a deterministic near-linear-time algorithm for weighted planar instances and explicitly relates Network Diversion to Two-Sets Cut-Uncut [16]. The relationship is useful for RPMEC because both problems combine separation and preservation requirements. However, RPMEC preserves reachability among protected sources, whereas Network Diversion preserves a surviving route through a specified bottleneck edge.

### Parameterized, exact, and declarative approaches

A second line of related work concerns exact methods for constrained graph separation. Parameterized algorithms for cut problems often exploit small terminal sets, small cut size, treewidth-like structure, or special graph classes; randomized contractions have been particularly influential for several parameterized cut problems [17]. These methods are complementary to the Dijkstra-style algorithm studied here. The Dijkstra-style algorithm is extremely efficient when its label-setting property holds, but it is not a general exact algorithm on arbitrary graphs. Parameterized or exponential-time methods, in contrast, can be used when no frontier-realizability certificate is available.

The ASP formulation in this paper belongs to the declarative optimization tradition. Answer Set Programming provides a compact way to encode combinatorial choices, recursive reachability, integrity constraints, and optimization objectives [10, 11]. Modern systems such as clingo combine grounding and solving in a single workflow and support incremental and optimization-oriented modeling [12]. For RPMEC, ASP is attractive because the two essential logical requirements are direct: *s*_2_ must remain reachable from *s*_1_, while *t* must not be reachable from *s*_1_ after the selected edges are cut. This makes ASP a useful exact fallback on general graphs, especially when the Dijkstra-style algorithm is not certified by planarity, laminarity, or dominator-compatible reachability.

### Biological pathway and interaction graphs

RPMEC has natural biological interpretations: one may want to block reachability to an undesirable state, disease module, toxic metabolite, or adversarial perturbation while preserving reachability among essential biological functions. Curated pathway resources provide several graph representations for such tasks. KEGG PATHWAY stores manually drawn maps of molecular interaction, reaction, and relation networks [5, 18]. Reactome provides a curated, peer-reviewed pathway knowledgebase with both hierarchical pathway organization and reaction-level diagrams [6, 19]. These curated resources are especially relevant to the Dijkstra-style algorithm when pathways are analyzed at the level of modules, subpathways, or planar pathway maps rather than at the level of all molecular interactions.

Protein and functional interaction networks provide a more general and often more difficult setting. BioGRID curates protein, genetic, and chemical interactions [7], while STRING integrates known and predicted physical and functional associations and, in its recent versions, supports regulatory directionality information [8]. Network-biology studies show that cellular networks often contain hubs, modular organization, motifs, and small-world or scale-free structure [9, 20, 21]. These features are biologically meaningful, but they also warn against applying the Dijkstra-style RPMEC algorithm blindly: hubs and cross-talk can create overlapping optimal regions that are not exposed by a single frontier relaxation.

Metabolic networks are another important application area. Since reactions may be reversible and metabolites may participate in many reactions, raw metabolic graphs can be dense and cyclic. A useful preprocessing idea is to contract strongly connected reaction structures into metabolic building blocks, yielding a reduced metabolic directed acyclic graph [22]. Recent MetaDAG work develops this idea as a tool for generating and analyzing metabolic networks through SCC contraction and m-DAG construction [23]. For RPMEC, this is significant because SCC-contracted metabolic graphs may become Dijkstra-solvable when the quotient graph is arborescent, laminar, or dominator-compatible. However, DAG structure alone is not enough: as shown by the directed hardness construction in this paper, arbitrary acyclic directed graphs can still encode global set-cover-like choices.

### How this work fits with previous work

The contribution of this paper is structural rather than merely algorithmic. Prior planar algorithms show that the Dijkstra-style method is correct under planar noncrossing structure [2]. Recent cut-uncut results show that broader connectivity-preserving cut problems are hard in general but tractable under particular planar or parameterized restrictions [3, 4]. Biological network resources motivate practical instances where the source-side represents essential molecular function and the destination represents a bad or disease-related state [5–8]. Our analysis connects these themes by identifying a common sufficient-and-necessary label-setting principle: the Dijkstra-style algorithm is exact exactly when every next optimum source-side region is frontier-realizable, and in directed graphs also reachability-uncrossable.

### Undirected RPMEC: Source-Side Formulation

Let *G* = (*V, E*) be an undirected graph with positive edge weights *w* : *E* → ℤ_*>*0_, and let *s*_1_, *s*_2_, *t* ∈ *V* be distinct terminals. For a vertex set *X* ⊆ *V*, let

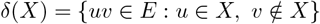

be the edge boundary of *X*, and let

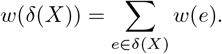

A source-side set *X* is feasible for the pair (*s*_1_, *v*) if

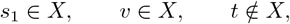

and *G*[*X*] is connected. Cutting all edges in *δ*(*X*) separates the connected protected component *X* from *t*. Therefore the true RPMEC value for connecting *s*_1_ to a vertex *v* while separating from *t* is

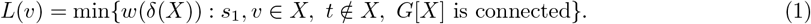

The desired three-terminal answer is *L*(*s*_2_).

For a set *A* ⊆ *V* with *s*_1_ ∈ *A* and *t* ∉ *A*, define

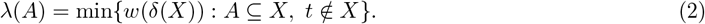

If *A* is connected, then any optimal solution to (2) may be chosen so that the source-side component containing *A* is connected. This is because all weights are positive: any disconnected component on the source-side that does not contain *A* can be moved to the *t*-side without increasing the cut value. Hence the operation in (2) is exactly the minimum cut operation used by the Dijkstra-style algorithm.

### The Undirected Dijkstra-Style Algorithm

The algorithm maintains a settled set *S* of vertices whose source-side components have been certified. For each settled vertex *s* ∈ *S*, let *C*_*s*_ denote the stored source-side component for *s*. Initially,

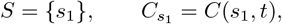

where *C*(*s*_1_, *t*) is the *s*_1_-side of a minimum *s*_1_–*t* cut.

At a generic step, the algorithm looks at every frontier edge *sv* ∈ *E* with

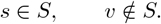

For each such pair (*s, v*), it computes

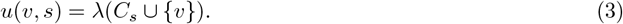

Here for any forced vertex set *A* ⊆ *V* with *t* ∉*A, λ*(*A*) denotes the value of the minimum edge cut separating *A* from *t*, obtained by forcing all vertices of *A* to lie on the source-side and computing a standard minimum *A*-to-*t* cut. Equivalently obtained by forcing all vertices in *A* to lie on the source-side and computing a standard minimum *A*-to-*t* cut. Equivalently, one may add a dummy source *D*, connect *D* to every vertex of *C*_*s*_ ∪ *{v}* with infinite-weight edges, and compute a minimum *D*–*t* cut. The source-side component of that cut is denoted 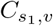. The algorithm chooses a pair (*s*^∗^, *v*^∗^) minimizing *u*(*v, s*), adds the resulting component to *S*, and assigns this value to newly reached vertices.

The analogy with Dijkstra’s shortest path algorithm is clear. In shortest paths, a shortest path to the next unsettled vertex must cross the frontier by one edge from an already settled vertex. Here, an optimum source-side cut region for the next unsettled vertex must be discoverable by forcing one frontier vertex *v* together with one already certified component *C*_*s*_.

### Exact Characterization in the Undirected Case

#### Definition 1

(Frontier-realizability). *Fix a tie-breaking rule for minimum cuts, so that every label L*(*v*) *has a canonical optimum source-side component C*_*v*_. *An algorithmic state S satisfies the* frontier-realizability condition *if*

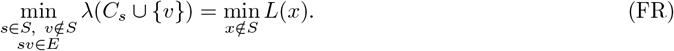

The left-hand side is the cheapest value visible to the Dijkstra-style algorithm through one frontier relaxation. The right-hand side is the cheapest true value among all unsettled vertices. Thus (FR) says that the next true optimum is visible from the current frontier.

#### Theorem 2

(Undirected frontier characterization). *For a fixed instance and a fixed tie-breaking rule, the Dijkstra-style algorithm is label-setting whenever every state before s*_2_ *is settled satisfies* (FR). *Conversely, if the algorithm is required to be label-setting at every state, then* (FR) *is necessary*.

*Proof*. We prove the sufficient direction first. Consider a state *S* satisfying (FR). For every frontier pair (*s, v*), the set computed in (3) is feasible for *v*, because it contains the already connected component *C*_*s*_, contains the neighbor *v*, and does not contain *t*. Therefore

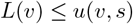

for every frontier pair. Hence no frontier candidate can be cheaper than the cheapest true unsettled label.

By (FR), however, at least one frontier candidate has value exactly equal to the cheapest true unsettled label. The algorithm chooses such a minimum candidate. Let *C* be the component it adds. Every newly added vertex *y* ∈ *C \ S* satisfies

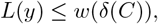

because *C* itself is feasible for *y*. If some newly added vertex had *L*(*y*) *< w*(*δ*(*C*)), then the right-hand side of (FR) would be smaller than the chosen candidate value, contradicting (FR). Hence every newly added vertex receives its true label. Repeating this argument proves label-setting correctness until *s*_2_ is settled.

For necessity, suppose the algorithm is label-setting at a state *S*. If (FR) failed, the cheapest frontier candidate would be strictly larger than min_*x*∉*S*_ *L*(*x*). The next vertex or component settled by the algorithm would then receive a label larger than some unsettled true label. This contradicts the label-setting property, which settles vertices in nondecreasing true value. Thus (FR) is necessary for label-setting correctness.

### Sufficient Graph and Instance Classes in the Undirected Case

The exact condition (FR) is state-dependent. It is therefore useful to identify structural conditions implying it.

#### Planar graphs

For undirected planar graphs, the correctness proof in [2] can be interpreted as showing that optimal source-side components are frontier-realizable. Intuitively, planar cut boundaries behave like noncrossing curves in the dual graph. If the cheapest unsettled optimal component were hidden behind another component in a way that no frontier relaxation could discover it, then two relevant cut boundaries would have to cross or create a hole. The planar no-crossing/no-hole argument rules out this obstruction.

Thus planar graphs are a clean sufficient class. This also includes planar subclasses such as outerplanar graphs, series-parallel graphs, trees, cactus graphs, and grid-like planar networks. The counterexample in Fig 1 shows that this planar argument does not extend to arbitrary graphs.

#### Laminar optimal source-side components

##### Definition 3

(Accessible laminar family). *The canonical optimum components {C*_*v*_ : *v* ∈ *V \ {t}} are* accessible laminar *if they are laminar by inclusion and, whenever C*_*x*_ *is not yet settled, there is a settled vertex s* ∈ *C*_*x*_ ∩ *S and a frontier edge sv with v* ∈ *C*_*x*_ *\ S such that*

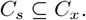

##### Corollary 4

(Laminar sufficient condition). *If the canonical optimum source-side components form an accessible laminar family, then the undirected Dijkstra-style algorithm is correct*.

*Proof*. Let *x* ∉*S* be an unsettled vertex of minimum true label *L*(*x*), and let *C*_*x*_ be its canonical optimum component. By accessibility, there is a frontier edge *sv* with *s* ∈ *S, v* ∈ *C*_*x*_ *\ S*, and *C*_*s*_ ⊆ *C*_*x*_. Since *C*_*x*_ contains *C*_*s*_ ∪ *{v}* and is feasible, the relaxation value satisfies

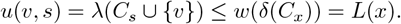

On the other hand, every frontier relaxation is feasible for its frontier vertex, so the minimum frontier value is at least min_*y*∉*S*_ *L*(*y*) = *L*(*x*). Therefore equality holds, which is exactly (FR).

This condition captures hierarchical or one-dimensional instances: rooted trees, path-like networks, nested ring networks, and weighted graphs where the optimum protected regions expand in a single inclusion order.

#### Cut-planar or noncrossing cut skeletons

Planarity of the entire graph is stronger than necessary. It is enough that the *relevant cut skeleton* is planar or noncrossing. More precisely, suppose that after contracting graph parts that always remain entirely on one side of every relevant optimum cut, the boundaries of all canonical optimum source-side components are noncrossing and hole-free. Then the same argument used in planar graphs implies frontier-realizability. This condition allows nonplanar subgraphs inside regions that are never split by the relevant cuts.

#### Rooted-greedoid viewpoint

The broadest abstraction is a rooted accessible set system. Let

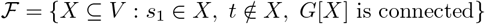

be the family of feasible source-side regions. The Dijkstra-style algorithm is correct when the optimum members of ℱ form an accessible family: every noninitial optimum region has a smaller optimum predecessor touching it by one frontier edge, and forcing that predecessor together with the frontier vertex does not increase the optimum value. This is exactly the set-system version of (FR).

### Why the Undirected Algorithm Fails on General Graphs

The failure mode in general graphs is not caused merely by cycles or high degree. It is caused by a hidden optimum component that requires simultaneous growth through two or more frontiers. In the counterexample of Fig 1, the globally optimal source-side component is

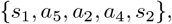

which combines vertices from both the left and right branches. A one-frontier relaxation from the already settled side instead discovers a left-chain or right-chain component of value 189. The hidden component of value 188 is not visible to the algorithm before it commits to the locally visible component. Planarity forbids this kind of interlocking through noncrossing cut boundaries; arbitrary graphs do not.

### Directed RPMEC

We now consider the directed analogue. Let *D* = (*V, A*) be a directed graph with positive arc weights *w* : *A* → ℤ_*>*0_. For *X* ⊆ *V*, define the outgoing cut

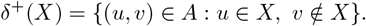

The directed three-terminal RPMEC asks for a minimum-cost set of arcs whose removal makes *t* unreachable from *s*_1_, while preserving reachability from *s*_1_ to *s*_2_. Since *s*_1_ ⇝ *s*_2_ is preserved, forbidding *s*_1_ ⇝ *t* also forbids *s*_2_ ⇝ *t*; otherwise *s*_1_ ⇝ *s*_2_ ⇝ *t*.

A source-side set *X* is feasible for vertex *v* if

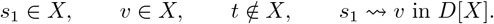

The true directed label is

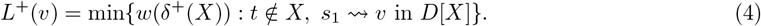

The desired value is *L*^+^(*s*_2_).

It is harmless to restrict attention to source-side sets in which every vertex is reachable from *s*_1_. Indeed, if *X* is feasible and *R* = Reach_*D*[*X*]_(*s*_1_), then *v* ∈ *R, t* ∉*R*, and there is no arc from *R* to *X \ R*. Therefore

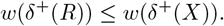

Thus an optimum solution can always be represented by an *s*_1_-reachable source-side region.

### Directed Dijkstra-Style Relaxation

In the directed algorithm, the frontier consists of outgoing arcs

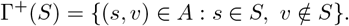

For a frontier arc (*s, v*), the algorithm computes

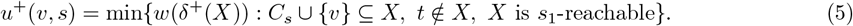

This can be implemented by a directed minimum cut with a dummy source *D*: add infinite-capacity arcs from *D* to *v* and to every vertex of *C*_*s*_, then compute a minimum *D*–*t* cut. If the returned source-side contains vertices not reachable from *s*_1_ inside that source-side, remove those unreachable vertices; by the argument above, this does not increase the outgoing cut value and does not remove any forced vertex in *C*_*s*_ ∪ *{v}*.

The directed case is harder than the undirected case because overlap of vertex sets is not enough. Two source-side regions may intersect, but their intersection may fail to contain a directed path from *s*_1_ to the predecessor vertex. This reachability failure is the directed analogue of crossing cut boundaries.

### Exact Characterization in the Directed Case

#### Definition 5

(Directed frontier-realizability). *A directed algorithmic state S satisfies the directed frontier-realizability condition if*

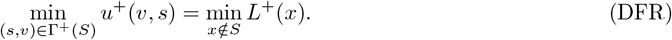

#### Theorem 6

(Directed frontier characterization). *For a fixed directed instance and tie-breaking rule, the directed Dijkstra-style algorithm is label-setting whenever every state before s*_2_ *is settled satisfies* (DFR). *Conversely*, (DFR) *is necessary if the algorithm is required to be label-setting at every state*.

*Proof*. The proof is the directed analogue of the undirected proof. Every relaxation value *u*^+^(*v, s*) is feasible for *v*, because *C*_*s*_ is reachable from *s*_1_, the arc (*s, v*) makes *v* reachable, and the source-side excludes *t*. Hence

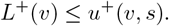

Therefore the cheapest frontier relaxation is never below the cheapest true unsettled label. If (DFR) holds, the cheapest frontier relaxation is exactly the cheapest true unsettled label. Any newly settled vertex has true value at most the relaxation value; if it had a strictly smaller true value, it would contradict the equality in (DFR). Hence the algorithm settles only correct labels.

The necessity argument is the same as before. If (DFR) failed at a state where label-setting correctness is required, the algorithm would settle a frontier candidate whose value is larger than the true value of some unsettled vertex, contradicting the label-setting property.

### Reachability-Uncrossing: A Useful Directed Sufficient Condition

The exact condition (DFR) is useful but state-dependent. The following theorem gives a more structural sufficient condition.

#### Theorem 7

(Directed reachable-uncrossing condition). *Assume the algorithm has correctly settled all vertices in S. Let x* ∉*S be an unsettled vertex minimizing L*^+^(*x*), *and let C*_*x*_ *be a canonical optimum source-side region for x. Suppose there is an s*_1_ ⇝ *x path P* ⊆ *C*_*x*_ *such that, if* (*s, v*) *is the first arc of P leaving S, then*

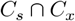

*contains an s*_1_ ⇝ *s path. Then the frontier relaxation through* (*s, v*) *has value at most L*^+^(*x*), *and therefore the state satisfies* (DFR).

*Proof*. Let

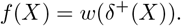

The directed cut function *f* is submodular:

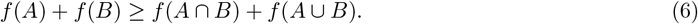

Because *C*_*s*_ ∩ *C*_*x*_ contains an *s*_1_ ⇝ *s* path and does not contain *t*, it is feasible for *s*. Since *C*_*s*_ is an optimum region for *s*, we have

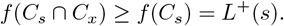

Also *f* (*C*_*x*_) = *L*^+^(*x*). Applying (6) with *A* = *C*_*s*_ and *B* = *C*_*x*_ gives

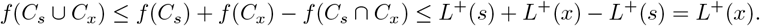

The set *C*_*s*_ ∪ *C*_*x*_ contains *C*_*s*_ and contains the frontier vertex *v*. It may include vertices not reachable from *s*_1_, but removing those unreachable vertices cannot increase the outgoing cut value. The reachable part still contains *C*_*s*_ and *v*, because *C*_*s*_ is reachable and (*s, v*) is an arc. Hence the relaxation value satisfies

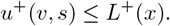

Since *x* was chosen with minimum true unsettled label, the frontier minimum cannot be below *L*^+^(*x*). Thus equality holds in (DFR).

### Directed Classes Where the Dijkstra-Style Algorithm Is Safe

The following classes are best understood as sufficient instance classes. They are not claimed to be a complete graph-minor classification.

#### Directed laminar optimum regions

If the canonical directed optimum regions *{C*_*v*_*}* are laminar by inclusion and accessible from *s*_1_, then the directed Dijkstra-style algorithm is correct. The proof is immediate from the previous theorem: if *C*_*s*_ ⊆ *C*_*x*_, then *C*_*s*_ ∩ *C*_*x*_ = *C*_*s*_, which certainly contains an *s*_1_ ⇝ *s* path.

#### Arborescent or unique-path reachability

Suppose the relevant source-reachable part of the directed graph has a rooted arborescent structure, meaning that each vertex has a unique relevant path from *s*_1_, or more generally that every feasible path to a vertex shares a fixed prefix. Then an optimum region for a later vertex must contain the same prefix used to reach any predecessor *s*. Therefore *C*_*s*_ ∩ *C*_*x*_ preserves the required *s*_1_ ⇝ *s* path, and the reachable-uncrossing theorem applies.

#### Dominator-compatible instances

A vertex *p* dominates a vertex *v* if every directed path from *s*_1_ to *v* passes through *p*. If each optimum region can be exposed through a frontier arc whose tail is a dominator of the remaining part of the optimum region, then the already settled region and the new optimum region share the necessary directed prefix. Hence the reachable-uncrossing condition holds.

#### Directed cut-planar skeletons with reachability compatibility

A directed planar drawing or a planar cut skeleton is not by itself the same as the undirected planar case, because orientations matter. The relevant sufficient condition is stronger: boundaries should be noncrossing, and intersections of optimum regions should preserve the directed path from *s*_1_ to the frontier predecessor. When this reachability-compatible noncrossing condition holds, the Dijkstra-style algorithm is safe by the reachable-uncrossing theorem.

#### Strongly connected block contractions

In some network instances, the graph decomposes into strongly connected blocks, and every relevant optimum region either contains a whole block or excludes it. If contracting those blocks produces a laminar, arborescent, or reachability-compatible cut skeleton, then the Dijkstra-style algorithm is correct on the original instance. The strongly connected blocks behave as atomic units, while the contracted skeleton supplies the required frontier-realizability.

### Acyclic Directed Graphs Remain NP-Complete

We consider the RPMEC problem in directed graph.

#### Decision problem

The directed DAG-RPMEC decision problem is the following. Given a weighted DAG *D* = (*V, A*), terminals *s*_1_, *s*_2_, *t*, and budget *B*, decide whether there is a set of arcs *F* ⊆ *A* with

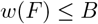

such that in *D* − *F*,

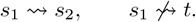

The problem is in NP because, after guessing *F*, both reachability conditions can be checked in linear time.

#### Reduction from Weighted Set Cover

We reduce from Weighted Set Cover [24]. Let the set-cover instance have universe

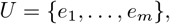

sets

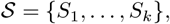

positive integer weights *a*_*j*_ for set *S*_*j*_, and budget *W*. Let

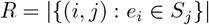

be the number of element–set incidences. Set

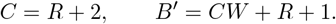

Let *M > B*^*′*^ be a prohibitive cost.

**Step 1: Element layers**. Create connector vertices

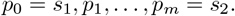

For each incidence *e*_*i*_ ∈ *S*_*j*_, create a branch

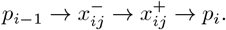

The middle arc 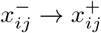 has cost 1. The two connector arcs have cost *M*.

**Step 2: Set-selection arcs**. For each set *S*_*j*_, create vertices 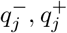 and a set-selection arc

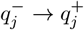

of cost *Ca*_*j*_.

**Step 3: Escape arcs to the destination**. For every incidence *e*_*i*_ ∈ *S*_*j*_, add a prohibitive arc

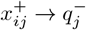

of cost *M*, and add a prohibitive arc

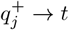

of cost *M*. Finally add the direct arc

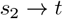

of cost 1.

The graph is acyclic. A topological order is obtained by listing the element layers in order,

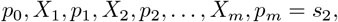

then the set-selection vertices 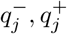, and finally *t*.

#### Completeness

Assume the Set Cover instance has a cover 𝒪 ⊆ 𝒮 with total weight at most *W*. For each element *e*_*i*_, choose one set *S*_*j*(*i*)_ ∈ 𝒪 that covers it. In the RPMEC instance, keep the branch corresponding to *S*_*j*(*i*)_ in layer *i*, and cut the middle arc of every other branch. Also cut the set-selection arc 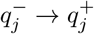 for every *S*_*j*_ ∈ 𝒪, and cut the direct arc *s*_2_ → *t*.

The remaining graph still has a path

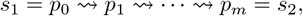

because one branch is kept in every element layer. Every kept branch points only to a selected set vertex 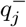, and the selected set-selection arc has been cut. The direct arc *s*_2_ → *t* has also been cut. Thus *s*_1_ cannot reach *t*. The total cost is at most

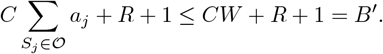

Therefore a set cover gives a feasible directed RPMEC cut.

#### Soundness

Conversely, suppose there is a feasible RPMEC cut *F* of cost at most *B*^*′*^. No prohibitive arc of cost *M* can be cut, since *M > B*^*′*^. Because *s*_1_ must still reach *s*_2_, each element layer must contain at least one uncut branch. If the surviving branch for element *e*_*i*_ corresponds to set *S*_*j*_, then 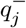 is reachable from *s*_1_ through the arc 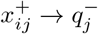. Since the arc 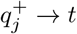 is prohibitive and cannot be cut, the only way to prevent reachability to *t* is to cut the set-selection arc

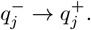

Thus, for every element *e*_*i*_, at least one set containing *e*_*i*_ has its set-selection arc cut. The cut set therefore defines a set cover.

It remains to check the budget. If the selected sets had total weight at least *W* + 1, then their set-selection arcs alone would cost at least

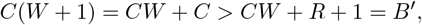

because *C* = *R* + 2. This contradicts the assumed budget. Hence the selected sets have total weight at most *W*.

##### Theorem 8.

*Directed edge-deletion RPMEC is NP-complete on DAGs*.

#### Hardness of Acyclic Graph Cases

Acyclicity alone does not imply directed frontier-realizability. In fact, the directed edge-deletion RPMEC decision problem remains NP-complete even when the input directed graph is a DAG, shown as follows.

*Proof*. Membership in NP follows from polynomial-time reachability checking. The construction above is polynomial and maps a Weighted Set Cover instance to a DAG-RPMEC instance. Completeness and soundness show that the Set Cover instance has a cover of weight at most *W* if and only if the DAG-RPMEC instance has a feasible cut of cost at most *B*^*′*^. Therefore DAG-RPMEC is NP-hard, and hence NP-complete.

The DAG hardness result has an important algorithmic consequence. Acyclicity alone cannot be a sufficient condition for the Dijkstra-style algorithm to be optimal on all directed instances, unless *P* = *NP*. A DAG may still contain many incompatible *s*_1_ ⇝ *s*_2_ route choices. The cost of cutting all escapes to *t* depends on a global set-cover-like choice across all layers, not on a locally visible one-frontier relaxation.

Therefore the correct structural condition remains frontier-realizability:

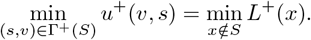

Acyclicity does not imply this equality. Laminarity, arborescent reachability, dominator compatibility, or reachability-compatible noncrossing cut skeletons are sufficient because they force the next optimum region to be exposed through one outgoing frontier arc from an already certified region.

### Open Complexity of the Three-Terminal Edge Case and Exact Algorithms

This section discusses the complexity status of the original three-terminal undirected edge version of RPMEC. The input is a connected undirected graph *G* = (*V, E*) with positive edge weights, two protected terminals *s*_1_, *s*_2_, and a destination terminal *t*. The goal is

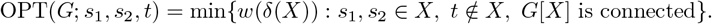

The work in [2] proved hardness for the node-cut version and gave a polynomial-time algorithm for the edge-cut version on planar graphs, but they explicitly left the complexity of CPMEC/RPMEC on general graphs open. The counterexample discussed in the introduction shows that the Dijkstra-style planar algorithm does not extend directly to arbitrary graphs, but it does not by itself prove NP-hardness. Thus the general three-terminal edge case remains a natural target for either a polynomial-time algorithm or a hardness proof.

The discussion below has two purposes. First, it records several reductions and proof strategies that clarify why the open case is subtle. Second, it gives exact algorithms that are useful in practice even without resolving the polynomial-time versus NP-hardness question.

#### A bond view of optimal solutions

The source-side formulation has an important consequence: with positive edge weights, an optimal solution can be assumed to define a bond-like cut.

##### Lemma 9

(Connected target side). *Let X be an optimal source-side set for three-terminal RPMEC. Then G*[*X*] *is connected by feasibility, and G*[*V \ X*] *may be assumed connected as well*.

*Proof*. Let *Y* be a connected component of *G*[*V \ X*] that does not contain *t*. Since *Y* has no vertex *t*, moving *Y* from the target side into the source-side preserves the condition *t* ∉*X*. Since *G* is connected, *Y* has at least one edge to *V \ Y*. Because *Y* is a connected component of *G*[*V \ X*], no such edge can go to (*V \ X*) *\ Y*. Hence *Y* has an edge to *X*, so *G*[*X* ∪ *Y*] remains connected. Moreover, all boundary edges between *X* and *Y* disappear from the cut. Since all edge weights are positive, the cut value strictly decreases. Hence no optimal solution has such a component *Y*. Therefore every component of *G*[*V \ X*] contains *t*, so the target side is connected.

Lemma 9 shows that the original three-terminal edge problem is a one-sided cut-uncut problem: the source-side must contain *s*_1_ and *s*_2_ in one connected component, while the target side has only one prescribed terminal *t* and can be taken connected automatically. This explains why the problem is close to Two-Sets Cut-Uncut, but also why it is not identical to the four-terminal case where two prescribed terminals must remain connected on the target side. Two-Sets Cut-Uncut asks for a cut separating two terminal sets *S* and *T* while preserving connectivity inside each set [3, 4]. Recent work shows that the general cut-uncut family is computationally challenging, while planar and parameterized cases admit specialized algorithms [3, 4, 25].

#### Relation to Network Diversion

The three-terminal RPMEC problem is closely related to Network Diversion. In Network Diversion, one is given terminals *s, t* and a designated edge *b*, and the goal is to remove a minimum set of edges so that every remaining *s*-*t* path uses *b*, while at least one *s*-*t* path remains. Directed Network Diversion is NP-hard, while the complexity of undirected Network Diversion on general graphs remains open; recent work gives efficient algorithms for planar Network Diversion [16].

There is a simple reduction from three-terminal RPMEC to undirected Network Diversion. Given an RPMEC instance (*G, s*_1_, *s*_2_, *t*), add a new edge

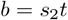

with very large weight or with the rule that *b* cannot be deleted. Set the Network Diversion terminals to *s* = *s*_1_ and *t* = *t*, with designated edge *b*. If *F* is an RPMEC solution in *G*, then in (*G* + *b*) − *F* the terminals *s*_1_ and *t* are connected through the path from *s*_1_ to *s*_2_ followed by *b*, and every *s*_1_-*t* path must use *b*, because *F* separates *s*_1_ from *t* in *G*. Conversely, if *F* is a Network Diversion solution for (*G* + *b, s*_1_, *t, b*), then after deleting *b* there is no *s*_1_-*t* path, while the remaining *s*_1_-*t* path using *b* implies that *s*_1_ reaches *s*_2_ in *G* − *F*. Hence *F* is an RPMEC solution.

This reduction is useful but it is in the “algorithmic” direction: a polynomial-time algorithm for undirected Network Diversion would imply a polynomial-time algorithm for three-terminal RPMEC. It does not prove hardness of RPMEC, because the open undirected Network Diversion problem is at least as broad as the constructed diversion instance. This is not evidence that RPMEC is harder than Network Diversion; rather, it shows that RPMEC is no harder than undirected Network Diversion under this transformation.

#### A conditional hardness route: pinned target-pair cut-uncut

A tempting way to prove NP-hardness is to reduce from the four-terminal case of Two-Sets Cut-Uncut with *S* = *{a, b}* and *T* = *{c, d}*. The difficulty is that RPMEC has only one target terminal. A natural pinning trick is to add a very expensive edge *cd* and take *t* = *c*, so that any low-cost RPMEC solution must place *d* on the *t*-side. This gives a correct reduction from the following restricted problem.

##### Definition 10

(Pinned target-pair cut-uncut). *In the pinned target-pair variant, the input is a weighted graph with terminals a, b, c, d, a budget B, and an uncuttable or weight-M > B edge cd. The task is to delete edges of total weight at most B so that a and b remain connected, a, b are separated from c, and the edge cd is not deleted. Equivalently, d is pinned to the c-side by an uncuttable edge*.

##### Lemma 11

(Pinned-pair reduction). *If the pinned target-pair cut-uncut problem is NP-hard, then weighted three-terminal RPMEC is NP-hard*.

*Proof*. Given a pinned instance with terminals *a, b, c, d*, create a three-terminal RPMEC instance on the same graph by setting

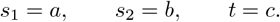

The uncuttable edge *cd* is kept in the graph. Any feasible pinned solution is immediately feasible for RPMEC, since it keeps *a, b* connected and separates them from *c*. Conversely, any RPMEC solution of cost at most *B* cannot cut *cd*. If *d* were placed on the source-side with *a, b*, then the uncut edge *cd* would connect the source-side to *t* = *c*, contradicting separation. Hence *d* is forced onto the *c*-side. Thus the same edge set is a feasible pinned solution. The two instances have the same optimum value.

Lemma 11 is not a full hardness proof, because the pinned target-pair problem itself must still be shown hard. Its value is that it isolates a promising target: proving hardness even when the second target terminal is pinned to *t* by an uncuttable edge would immediately resolve the weighted three-terminal RPMEC case.

#### Why a direct Set Cover reduction is difficult

The most natural NP-hardness attempt is a reduction from Set Cover. For each element *e*_*i*_, one creates a layer between consecutive connector vertices

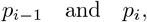

where *p*_0_ = *s*_1_ and *p*_*m*_ = *s*_2_. For every set *S*_*j*_ containing *e*_*i*_, one creates an occurrence branch through a vertex *x*_*ij*_ :

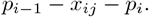

Choosing a source-side path from *s*_1_ to *s*_2_ then resembles choosing one set occurrence for every element. To charge for selected sets, one adds a set vertex *q*_*j*_ and connects every occurrence *x*_*ij*_ with *e*_*i*_ ∈ *S*_*j*_ to *q*_*j*_. The intended behavior is:

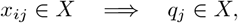

and placing *q*_*j*_ on the source-side incurs the set-selection cost through an edge from *q*_*j*_ toward *t*.

The obstacle is that the graph is undirected. Once *q*_*j*_ is adjacent to many occurrences, it can create unintended shortcuts:

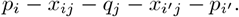

Such shortcuts may allow the preserved *s*_1_-*s*_2_ connection to skip layers, so the construction no longer forces one valid choice for every element. This is precisely why the directed version is easier to prove hard: arc directions can allow membership implications without allowing reverse shortcuts. In the undirected three-terminal edge problem, a successful Set Cover proof likely needs a *no-shortcut membership gadget* : a gadget that forces *q*_*j*_ to be paid for when an occurrence *x*_*ij*_ is used, but does not let *q*_*j*_ act as a connector between unrelated element layers.

Fig 3 illustrates the desired construction and the shortcut that must be blocked.

**Fig 3.**
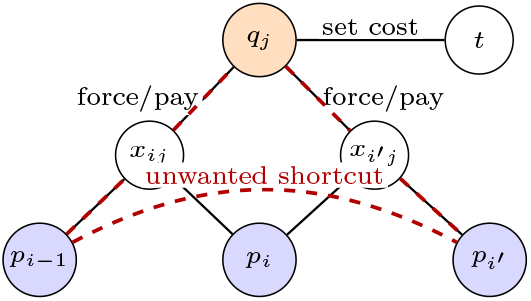
A Set Cover hardness attempt and its obstacle. The set vertex *q*_*j*_ should charge the solution when an occurrence *x*_*ij*_ is used. In an undirected graph, however, *q*_*j*_ may also connect different occurrence layers and create shortcuts in the preserved *s*_1_-*s*_2_ path.

#### A hard layer-respecting variant

Although the unrestricted Set Cover reduction has the shortcut problem above, it becomes valid under a natural extra assumption. Define the *layer-respecting RPMEC* problem by requiring that every feasible preserved *s*_1_-*s*_2_ connection pass through the element layers in order, and by forbidding set-selection vertices from being used as connectors between layers. This restriction can model settings where layers represent time, pathway stage, or policy zones and off-layer traversal is disallowed.

##### Theorem 12

(Conditional hardness under layer-respecting structure). *The layer-respecting three-terminal RPMEC problem is NP-hard, even with positive edge weights*.

*Proof sketch*. Reduce from Weighted Set Cover. Let *U* = *{e*_1_, …, *e*_*m*_*}* be the universe, let 𝒮 = *{S*_1_, …, *S*_*r*_*}* be the family of sets with weights *w*_*j*_, and let *W* be the set-cover budget. Create connectors *p*_0_ = *s*_1_, *p*_1_, …, *p*_*m*_ = *s*_2_. For each occurrence *e*_*i*_ ∈ *S*_*j*_, create a branch *p*_*i*−1_ − *x*_*ij*_ − *p*_*i*_. A layer-respecting source-side connection from *s*_1_ to *s*_2_ must choose at least one branch in each element layer. For each set *S*_*j*_, create a set vertex *q*_*j*_. The construction enforces that if any occurrence *x*_*ij*_ is used on the source-side, then *q*_*j*_ must also be on the source-side; otherwise a prohibitive edge would be cut. Placing *q*_*j*_ on the source-side cuts a set-cost edge of weight proportional to *w*_*j*_.

If there is a set cover of weight at most *W*, choose one occurrence branch per element covered by the selected sets, place the corresponding set vertices on the source-side, and cut the set-cost edges for the selected sets. This yields a feasible RPMEC solution of the corresponding cost. Conversely, any low-cost feasible RPMEC solution must contain a layer-respecting path from *s*_1_ to *s*_2_, hence at least one occurrence from every element layer. The forced set vertices corresponding to these occurrences form a set cover. By scaling the set-cost edges above the total possible branch-cleanup cost, the selected sets have total weight at most *W* if and only if the RPMEC solution has cost at most the constructed budget.

Theorem 12 should be read as a hardness guide rather than a resolution of the open case. It identifies the exact missing ingredient for an unrestricted reduction: a purely undirected gadget that enforces the occurrence-to-set payment implication without permitting shortcuts through set vertices.

#### An exact path-cut formulation

The following identity is useful both algorithmically and conceptually. Let 𝒫 (*s*_1_, *s*_2_) be the set of simple *s*_1_-*s*_2_ paths. For a path *P*, define

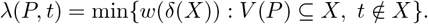

This value is computed by one ordinary minimum cut after forcing all vertices of *P* onto the source-side, equivalently by adding a dummy source adjacent to every vertex of *P* with infinite-weight edges.

##### Lemma 13

(Path-cut identity). *For three-terminal RPMEC*,

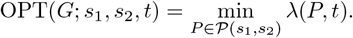

*Proof*. Let *X* be any feasible RPMEC source-side. Since *G*[*X*] is connected and contains *s*_1_, *s*_2_, it contains an *s*_1_-*s*_2_ path *P*. Because *X* is feasible for the minimization defining *λ*(*P, t*), we have *λ*(*P, t*) ≤ *w*(*δ*(*X*)).

Taking *X* to be an optimum source-side gives

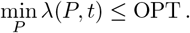

Conversely, for any path *P*, every set *X* feasible for *λ*(*P, t*) contains all vertices of *P*, and therefore contains a connected *s*_1_-*s*_2_ subgraph. Thus such an *X* is feasible for RPMEC, so OPT ≤ *λ*(*P, t*) for every *P*. Taking the minimum over *P* gives the reverse inequality.

Lemma 13 explains both the promise and the difficulty of the problem. For a fixed path, the optimum cut is polynomial-time computable. The challenge is that there may be exponentially many *s*_1_-*s*_2_ paths, and the value *λ*(*P, t*) is not additive along the path. This non-additivity is exactly what breaks a one-label Dijkstra-style dynamic program in general graphs.

#### FPT algorithm by cut size

The open polynomial-time status does not prevent useful exact parameterized algorithms. For unweighted graphs, three-terminal RPMEC is fixed-parameter tractable parameterized by the optimum cut size *k*.

##### Theorem 14

(FPT by important edge separators). *The unweighted three-terminal RPMEC decision problem can be solved in* 4^*k*^*n*^*O*(1)^ *time, where k is the allowed cut size*.

*Proof*. Enumerate all important (*s*_1_, *t*) edge separators of size at most *k*. The standard important-separator theorem gives at most 4^*k*^ important separators of size at most *k*, and they can be enumerated within the same bound up to polynomial factors; the edge version follows from the usual conversion of edge cuts to vertex cuts by subdividing edges [26, 27].

For each enumerated separator *F*, compute the connected component *R*_*F*_ of *s*_1_ in *G* − *F*. Accept if *s*_2_ ∈ *R*_*F*_ and *t* ∉*R*_*F*_.

To prove correctness, suppose there is a feasible RPMEC cut *F* of size at most *k*. Let *R* be the component of *s*_1_ in *G* − *F*. Then *s*_2_ ∈ *R, t* ∉*R*, and *δ*(*R*) is an (*s*_1_, *t*) edge separator of size at most |*F* | ≤ *k*. By the domination property of important separators, there is an important (*s*_1_, *t*) separator *F* ^*′*^ of size at most |*δ*(*R*)| whose *s*_1_-reachable side *R*^*′*^ contains *R*. Since *s*_2_ ∈ *R* ⊆ *R*^*′*^ and *t* ∉*R*^*′*^, the separator *F* ^*′*^ is feasible for RPMEC. Therefore the enumeration will find a feasible cut whenever one exists.

For integer-weighted instances, one obtains a pseudo-polynomial variant by replacing an edge of weight *c* with *c* parallel unit edges or an equivalent unit-capacity gadget, so the parameter becomes the optimum total cost. For arbitrary large weights, the ASP/IP formulations below are usually more appropriate.

#### Multi-label exact search over closed source regions

The Dijkstra-style algorithm fails because it stores only one label per vertex. A direct exact repair is to store multiple source-side regions. For any connected set *A* with *s*_1_ ∈ *A* and *t* ∉*A*, define its cut closure

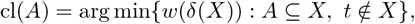

using a canonical tie-breaking rule. This closure is found by an ordinary minimum cut with *A* forced to the source-side.

The exact multi-label algorithm is a uniform-cost search over closed regions.

1. Start with *C*_0_ = cl(*{s*_1_*}*).
2. Store closed regions in a priority queue keyed by *w*(*δ*(*C*)).
3. Pop the minimum-key region *C*. If *s*_2_ ∈ *C*, return *C*.
4. For every boundary edge *uv* with *u* ∈ *C* and *v* ∉*C*, generate

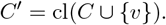
5. Insert *C*^*′*^ unless it is dominated by a previously generated region.

A simple dominance rule is

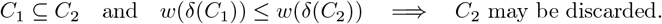

The algorithm is exact because every feasible source-side can be reached by adding the vertices of an internal *s*_1_-*s*_2_ path one at a time and closing after each addition. It may be exponential in the number of nondominated closed regions, but it is a principled exact version of the Dijkstra idea and can be effective when the number of relevant closed regions is small.

#### ASP and integer-programming formulations

For weighted practical instances, declarative optimization is a robust fallback. The ASP formulation in the present draft chooses cut edges, defines reachability from *s*_1_ through non-cut edges, enforces that *s*_2_ remains reachable, forbids *t* from being reachable, and minimizes total cut cost. This directly expresses the RPMEC constraints and avoids relying on frontier-realizability.

A standard mixed-integer programming formulation is also possible. Use binary variables *x*_*v*_ indicating whether a vertex is on the source-side and *y*_*uv*_ indicating whether edge *uv* is cut. Add

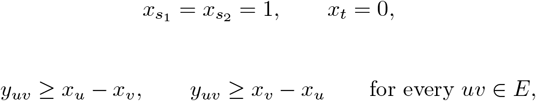

then minimize ∑_*uv*∈*E*_ *w*_*uv*_*y*_*uv*_. To enforce connectivity of *s*_1_ and *s*_2_ inside the source-side, add a one-unit flow from *s*_1_ to *s*_2_ using only vertices with *x*_*v*_ = 1. For each undirected edge *uv*, introduce directed flow variables *f*_*uv*_ and *f*_*vu*_ and impose capacity constraints such as

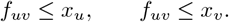

Flow conservation sends one unit from *s*_1_ to *s*_2_ and zero net flow through all other vertices. The resulting MILP is exact and naturally supports additional domain constraints such as forbidden cuts, policy zones, or uncertainty penalties.

#### Bounded-treewidth dynamic programming

Another exact algorithm is dynamic programming over a tree decomposition. For each bag, keep:

1. the side assignment of each bag vertex, source-side or target side;
2. the connectivity partition of source-side bag vertices;
3. optionally, the connectivity partition of target-side bag vertices, if one wants to enforce the bond form explicitly;
4. the accumulated cost of cut edges whose endpoints have already been introduced and forgotten.

At the root, accept only states in which *s*_1_ and *s*_2_ are in the same source-side component and *t* is on the target side. The number of connectivity partitions of a bag of size *τ* + 1 is Bell-number sized, so this gives a running time of the form

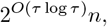

where *τ* is the treewidth. This is useful for pathway graphs, infrastructure graphs, and cyber graphs with moderate decomposability. It also complements the parameterized cut-size algorithm: treewidth DP is strong when the graph is structurally narrow, while important-separator enumeration is strong when the optimum cut is small.

#### Summary of hardness directions and algorithms

The current status can be summarized as follows.

| Approach | What it gives | Status |
| --- | --- | --- |
| Planar Dijkstra-style algorithm | Polynomial-time exact algorithm for planar CPMEC/RPMEC | Known positive result [2] |
| General Dijkstra-style algorithm | Fast greedy dynamic program based on one-frontier relaxations | Not exact in general; counterexample gives 189 versus 188 |
| Reduction to Network Diversion | RPMEC reduces to undirected Network Diversion by adding a designated edge $s_2t$ | Useful relationship, but not a hardness proof because undirected Network Diversion is open [16] |
| Pinned target-pair route | If pinned target-pair cut-uncut is NP-hard, then weighted three-terminal RPMEC is NP-hard | Conditional hardness target |
| Set Cover route | Would prove NP-hardness if a no-shortcut membership gadget is found | Main obstruction is undirected shortcutting through set vertices |
| Layer-respecting variant | Set Cover reduction works when layer shortcuts are explicitly forbidden | NP-hard restricted/assumption-based variant |
| Important separators | Enumerate important $(s_1, t)$ separators and test whether $s_2$ remains reachable | Exact FPT algorithm in $4^k n^{O(1)}$ time for unweighted cut size $k$ |
| Multi-label closed-region search | Uniform-cost search over closed source regions instead of one label per vertex | Exact exponential algorithm; often practical with strong dominance pruning |
| ASP/MILP | Directly encodes separation, preservation, and cost minimization | Exact practical fallback for weighted instances |
| Treewidth DP | Dynamic programming over side assignments and connectivity partitions | Exact in $2^{O(\tau \log \tau)} n$ time for treewidth $\tau$ |

Thus, the most defensible current position is that the general three-terminal undirected edge case remains open, but it is surrounded by strong structure. A polynomial-time algorithm would likely need to exploit a source-side closure structure stronger than the failed one-label Dijkstra rule. A hardness proof likely needs either a pinned target-pair construction or an undirected no-shortcut membership gadget. Meanwhile, important-separator enumeration, multi-label source-region search, ASP/MILP, and bounded-treewidth dynamic programming provide exact algorithms for substantial practical subclasses.

### Approximation Algorithms for the Three-Terminal Edge Case

The preceding section treats the open complexity of the original three-terminal undirected edge case. In this section we discuss approximation algorithms and approximable graph classes for the same problem. Throughout this section the input is an undirected graph *G* = (*V, E*) with positive edge weights *w* : *E* → ℝ_*>*0_ and terminals *s*_1_, *s*_2_, *t*. The objective is

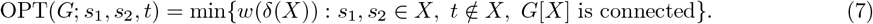

Here *δ*(*X*) denotes the set of edges with exactly one endpoint in *X*. Since the complexity of the general weighted problem remains open, the goal of this section is not to claim a universal constant-factor approximation for all graphs. Instead, we identify several approximation principles that are useful in structured instances and in practice:

1. a path-closure identity that converts RPMEC into a minimum over protected *s*_1_*s*_2_ paths;
2. an a posteriori lower bound that certifies the quality of any feasible solution;
3. exact algorithms for tractable graph families;
4. provable approximation guarantees for low-exposure, relaxed-cut-repairable, and near-tractable-core graphs.

The guiding message is that approximation becomes effective when the preserved *s*_1_–*s*_2_ connection has small exposure to *t*, or when the instance is close to a class on which RPMEC is exactly solvable.

#### A path-closure identity

Let *P* be an *s*_1_–*s*_2_ path in *G* − *t*. Define the *closure value* of *P* with respect to *t* by

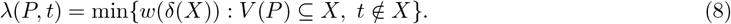

The value *λ*(*P, t*) is computable by one ordinary minimum cut computation: add a dummy source *d*, connect *d* to every vertex of *P* by an infinite-weight edge, and compute a minimum *d*–*t* cut. Let *X*_*P*_ be the source-side returned by this cut. The connected component of *G*[*X*_*P*_] containing *P* is a feasible RPMEC source-side, and removing all other source-side components cannot increase the boundary cost. Thus every path closure gives a feasible RPMEC solution.

##### Lemma 15

(Path-closure identity). *For the undirected three-terminal edge RPMEC problem*,

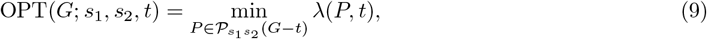

*where* 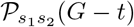 *is the set of simple s*_1_*–s*_2_ *paths in G* − *t*.

*Proof*. Let *X*^⋆^ be an optimum RPMEC source-side. Since *G*[*X*^⋆^] is connected and contains *s*_1_ and *s*_2_, it contains an *s*_1_–*s*_2_ path *P* ^⋆^. The set *X*^⋆^ is feasible for the cut problem defining *λ*(*P* ^⋆^, *t*), so

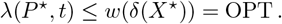

This proves that the right-hand side of (9) is at most OPT.

Conversely, for any path *P*, a source-side attaining *λ*(*P, t*) contains *V* (*P*) and excludes *t*. Taking the connected component containing *P* gives a connected set containing *s*_1_ and *s*_2_ and excluding *t*, with boundary weight no larger than the original source-side. Therefore *λ*(*P, t*) is at least OPT for every *P*, and equality follows.

Fig 4 illustrates the identity. The path *P* is forced to remain on the protected source-side. The rest of the source-side is then chosen by an ordinary minimum cut that separates the forced path from *t*.

**Fig 4.**
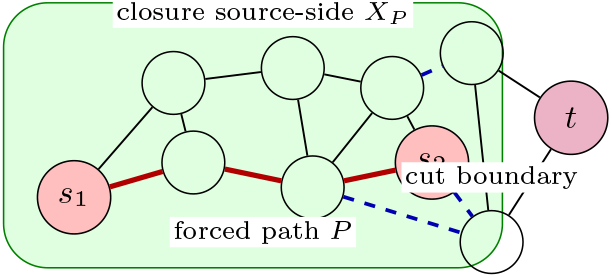
Path closure. A candidate protected path *P* from *s*_1_ to *s*_2_ is forced onto the source-side; an ordinary minimum cut then computes the cheapest source-side closure separating *P* from *t*.

**Fig 5.**
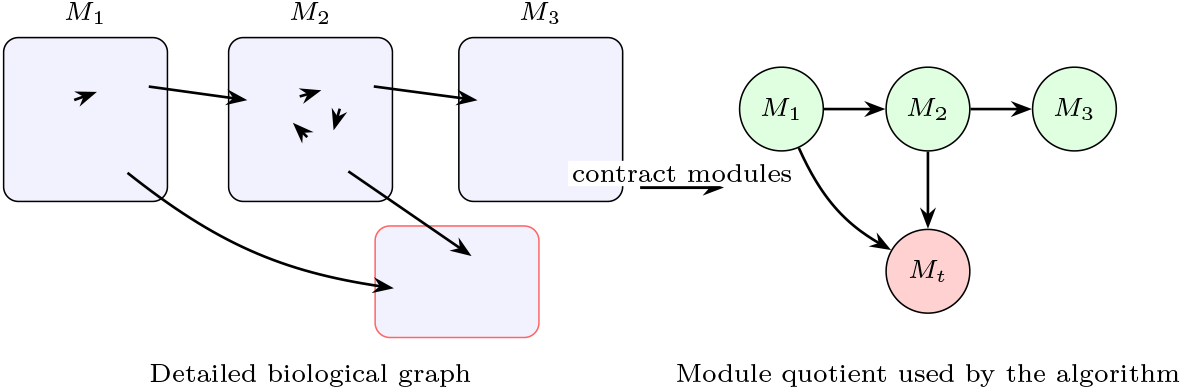
Figure. A common biological workflow: contract protein complexes, feedback SCCs, reversible metabolic blocks, or curated subpathways, then solve RPMEC on the quotient. The Dijkstra-style algorithm is exact when the quotient has a frontier-realizable structure, such as a tree, planar graph, out-arborescence, or dominator-compatible DAG.

#### A lower bound and a posteriori approximation certificates

A useful relaxation is obtained by forcing only the two terminals *s*_1_ and *s*_2_ to be on the source-side, without requiring this source-side to be connected. Define

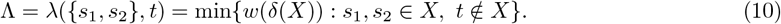

This is an ordinary minimum cut with a dummy source connected to *s*_1_ and *s*_2_ by infinite-weight edges. Since every feasible RPMEC source-side is feasible for this relaxation,

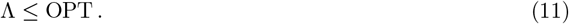

Consequently, any feasible RPMEC solution *X* has a certified a posteriori approximation ratio

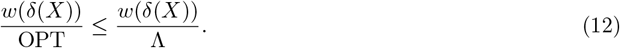

This certificate is important in practice. It means that a heuristic, a Dijkstra-style run, a path-closure algorithm, an ASP solver stopped early, or an MILP incumbent can all be accompanied by a rigorous instance-specific quality bound. If a solution has cost 105 and Λ = 100, then the solution is certified to be within a factor 1.05 of optimum, regardless of how it was found.

If the source-side of the relaxed cut attaining Λ is already connected, then the relaxation is tight and the relaxed solution is optimal for RPMEC. Thus many easy instances can be recognized by a single ordinary minimum cut.

#### Certified path-and-repair approximation

The path-closure identity suggests the following practical approximation algorithm. The algorithm searches over a selected family of promising *s*_1_–*s*_2_ paths, computes the exact closure value of each path, and returns the best closure. The quality is certified by the lower bound Λ.

This is a meta-algorithm rather than a universal constant-factor approximation. Its guarantee depends on the candidate path family.

##### Theorem 16

(Path-family approximation). *Let* ℱ *be a family of s*_1_*–s*_2_ *paths in G* − *t. If there exists P* ∈ ℱ *such that*

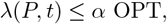

*then Algorithm 1 returns an α-approximation. Moreover, the returned ratio is always certified by* min_*P* ∈ℱ_ *λ*(*P, t*)*/*Λ.

*Proof*. The algorithm returns the path closure of minimum value among all paths in ℱ, so its cost is at most *λ*(*P, t*) ≤ *α* OPT. The certificate follows from Λ ≤ OPT.

##### Algorithm 1

Certified path-and-repair approximation for RPMEC

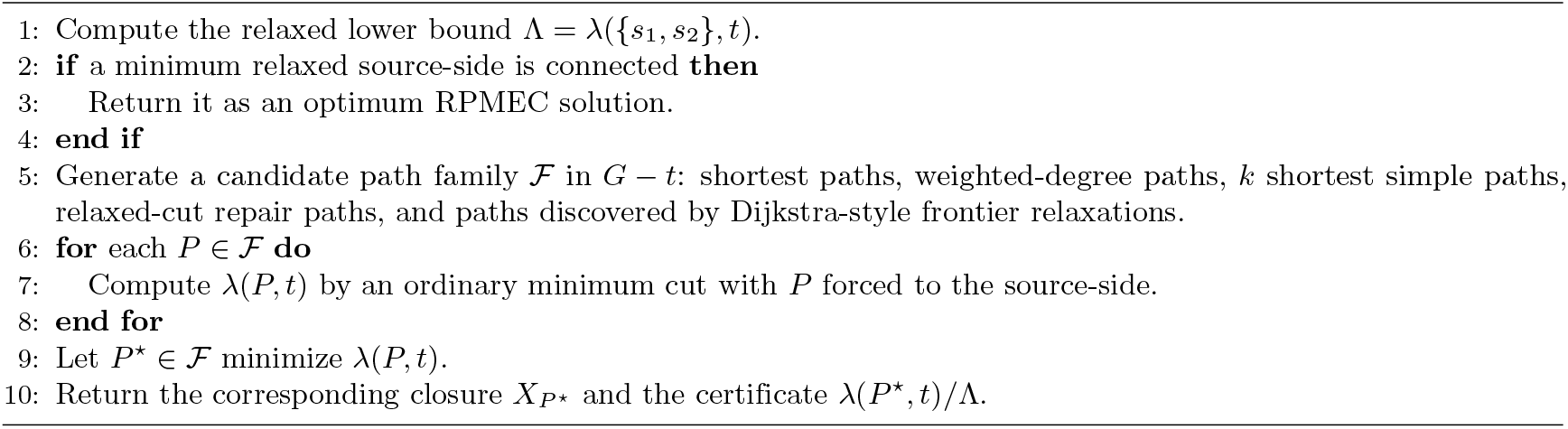

This theorem is deliberately simple, but it is useful: it separates the approximation problem into two parts. The min-cut closure is exact and polynomial-time; the only approximation question is whether the path generator finds a path whose closure is near-optimal.

#### Low-exposure path instances

For a vertex set *Y*, call *w*(*δ*(*Y*)) its *exposure*. For a path *P*, the exposure *w*(*δ*(*V* (*P*))) is an upper bound on the closure value:

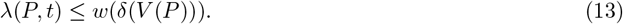

Therefore, if the graph contains an *s*_1_–*s*_2_ path whose exposure is small relative to optimum, then path closure gives a good approximation.

##### Definition 17

(Low-exposure path instance). *An RPMEC instance is α-path-exposed if there exists an s*_1_*–s*_2_ *path P in G* − *t such that*

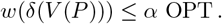

On such instances, any algorithm that finds such a path and closes it is an *α*-approximation. This class captures graphs in which the protected assets can be connected by a narrow corridor whose boundary is already close to an optimum isolating boundary.

A computable surrogate is obtained from weighted vertex degrees. Let

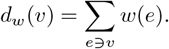

For every path *P*,

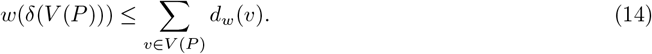

Thus one can compute a shortest *s*_1_–*s*_2_ path in *G* − *t* using vertex cost *d*_*w*_(*v*) and then close that path by a minimum cut. If the returned path is *P*, then the instance-specific guarantee is

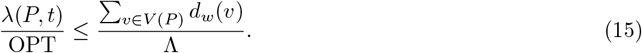

For unweighted graphs with maximum degree Δ and *s*_1_–*s*_2_ distance *d* in *G* − *t*, this gives

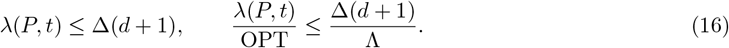

This bound is useful when protected terminals are close, degrees are bounded, and the relaxed terminal-to-target cut lower bound is not small.

#### Relaxed-cut-repairable graphs

Another approximation strategy starts from the relaxed lower-bound cut (10). Let *R* be a source-side attaining Λ. If *G*[*R*] is connected, then *R* is already optimal. Otherwise, let *R*_1_ and *R*_2_ be the connected components of *G*[*R*] containing *s*_1_ and *s*_2_, respectively. A repair path is a path *Q* in *G* − *t* that connects *R*_1_ to *R*_2_. Define the incremental repair cost of *Q* by

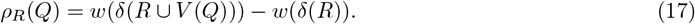

This value may be negative, but it is always an upper bound on the additional cost paid by adding the connector *Q* to the relaxed source-side.

##### Definition 18

(Repairable relaxed cut). *An instance is β-repairable if some minimum relaxed source-side R and some repair path Q satisfy*

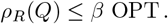

##### Theorem 19

(Relaxed-cut repair). *If an instance is β-repairable and the algorithm finds a repair path Q with ρ*_*R*_(*Q*) ≤ *β* OPT, *then closing R* ∪ *V* (*Q*) *gives a* (1 + *β*)*-approximation*.

*Proof*. The set *R* ∪ *V* (*Q*) contains *s*_1_ and *s*_2_, excludes *t*, and connects the two terminal-containing components of the relaxed source-side. Its closure has cost at most

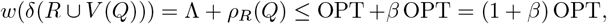

using Λ ≤ OPT.

This class is common in practice. The relaxed cut often already chooses the correct side of the target, but it may split the protected terminals into two nearby islands. If the islands can be connected through a low-exposure corridor, then a near-optimal RPMEC solution is obtained by repairing the relaxed cut.

#### Near-tractable-core approximation

Suppose that deleting a small exceptional edge set *R* ⊆ *E* turns *G* into a graph *H* = *G* − *R* from a class on which RPMEC can be solved exactly. Examples include trees, cactus graphs, outerplanar graphs, series-parallel graphs, planar graphs, bounded-treewidth graphs, or any graph satisfying the frontier-realizability condition from the frontier-realizability section. The following theorem gives an approximation guarantee when the exceptional edges have small total weight.

##### Theorem 20

(Approximation from an exact core). *Let H* = *G* − *R, where RPMEC can be solved exactly on H. Assume there exists an optimum RPMEC source-side X*^⋆^ *in G such that H*[*X*^⋆^] *is connected. Let X*_*H*_ *be an optimum RPMEC source-side in H. Then, when evaluated in the original graph G*,

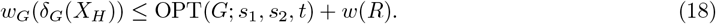

*Consequently, if w*(*R*) ≤ *ε* OPT, *then X*_*H*_ *is a* (1 + *ε*)*-approximation. If the stronger checkable condition w*(*R*) ≤ *ε*Λ *holds, then the same* (1 + *ε*) *guarantee follows*.

*Proof*. Because *H*[*X*^⋆^] is connected, *X*^⋆^ is feasible for RPMEC in *H*. Hence

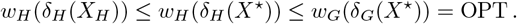

When *X*_*H*_ is evaluated in *G*, the only boundary edges not counted in *H* are edges of *R*. Thus

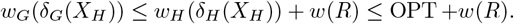

If *w*(*R*) ≤ *ε* OPT, the first ratio statement follows. The condition *w*(*R*) ≤ *ε*Λ is checkable and sufficient, because Λ ≤ *OPT* implies *w*(*R*) ≤ *εOPT*.

This theorem is useful for nearly planar, nearly tree-like, and modular graphs. For example, suppose a biological or cyber graph consists of a planar or tree-like backbone plus a small number of cross-talk edges. If the cross-talk edges have total weight at most 5% of the lower bound Λ, and deleting them does not destroy at least one optimum protected *s*_1_–*s*_2_ connection, then solving the backbone exactly and evaluating the resulting cut in the original graph gives a certified 1.05-approximation.

This statement is weaker than a general PTAS for minor-free or near-planar RPMEC, but it is robust and directly checkable. It is also aligned with approximation schemes for planar network problems, where a hard instance is often reduced to a structured core or spanner before dynamic programming is applied [28, 29]. In RPMEC, however, the protected-connectivity constraint must be preserved, so these results cannot be imported directly.

#### Exact special cases as approximation baselines

The approximation methods above are most useful when combined with exact algorithms for special graph classes.

##### Trees

If *G* is a tree, the unique *s*_1_–*s*_2_ path *P*_12_ must remain on the source-side. The optimum is exactly *λ*(*P*_12_, *t*), which in a tree is obtained by cutting the minimum-weight edge on the unique path from *t* to *P*_12_. Thus trees are solved exactly.

##### Polynomially many protected paths

If *G* − *t* has only polynomially many simple *s*_1_–*s*_2_ paths, then Lemma 15 yields a polynomial-time exact algorithm by enumerating all paths and computing *λ*(*P, t*) for each one. This includes many small cactus-like or corridor-like instances.

##### Planar and planar-subclass graphs

Planar RPMEC/CPMEC is polynomial-time solvable in the planar algorithm [2]. Therefore outerplanar graphs, cactus graphs, and series-parallel graphs are exact special cases as well. Later work on Two-Sets Cut-Uncut further supports the tractability of related cut-uncut problems on planar graphs [3].

##### Bounded treewidth

A standard connectivity dynamic program over a tree decomposition solves RPMEC exactly in time 2^*O*(*τ* log *τ*)^*n*, where *τ* is the treewidth. This gives exact algorithms for graphs whose global structure is narrow, even if they are not planar.

##### Small optimum cut

For unweighted graphs, the important-separator algorithm gives an exact fixed-parameter algorithm in the optimum cut size *k*. Important separators are a standard tool in parameterized cut algorithms [26, 27].

#### Relation to multiway-cut approximation

Classical Multiway Cut is a natural comparison point, but it is not a direct approximation algorithm for RPMEC. Multiway Cut with terminals *s*_1_, *s*_2_, *t* asks to separate all three terminals from one another. An RPMEC solution, in contrast, must keep *s*_1_ and *s*_2_ connected. Thus a multiway cut solution is generally infeasible for RPMEC.

This distinction is important because Multiway Cut has well-developed approximation theory. Dahlhaus, Johnson, Papadimitriou, Seymour, and Yannakakis gave the classical (2 − 2*/k*)-approximation for *k* terminals [30], and later work improved the approximation ratio via LP rounding [31]. Planar Multiway Cut also admits approximation schemes [29]. These results demonstrate that terminal-separation problems often have good approximations, but they do not automatically apply to cut-uncut problems such as RPMEC because the preservation constraint changes the feasible region.

The path-closure and repair algorithms above are therefore better aligned with the RPMEC structure: they first preserve an *s*_1_–*s*_2_ connection and then separate the resulting protected object from *t*.

#### Comparison of approximable graph classes

Table 3 summarizes the main approximation and exactness guarantees.

**Table 1.** Related algorithmic and biological work, organized by its connection to reachability-preserving minimum edge cuts.

| Area | Representative works | Main structure studied | Relation to RPMEC |
| --- | --- | --- | --- |
| Ordinary min cut | Ford–Fulkerson [1] | Two-terminal separation via flows | Provides the baseline; RPMEC adds preservation constraints that ordinary min cut does not model. |
| CPMC/RPMEC | Duan–Xu [2] | Connectivity-preserving node/edge cuts; planar CPMEC | Introduces the closest predecessor of RPMEC and the Dijkstra-style planar algorithm. |
| Two-Sets Cut-Uncut | Bentert et al. [3, 4] | Separate two terminal sets while preserving each set internally | Generalizes the cut-uncut viewpoint; explains why preservation constraints are algorithmically hard. |
| Network Diversion | Curet; Cullenbine et al.; Bentert et al. [14–16] | Force surviving paths through a designated edge | Related preservation/separation model; useful contrast to RPMEC. |
| ASP and exact modeling | Gebser et al.; Erdem et al. [10–12] | Declarative reachability and optimization constraints | Provides an exact fallback when Dijkstra-style frontier-realizability is not certified. |
| Curated pathways | KEGG, Reactome [5, 6, 18, 19] | Pathway maps, reaction diagrams, pathway hierarchies | Natural source of planar, hierarchical, or module-level RPMEC instances. |
| PPI and functional networks | BioGRID, STRING, network-biology studies [7–9] | Dense interaction graphs, hubs, modules, motifs | Motivates RPMEC applications but usually requires module contraction or certification before using Dijkstra. |
| Metabolic graph reductions | Metabolic building blocks and MetaDAG [22, 23] | SCC contraction and metabolic DAGs | Can produce arborescent or dominator-compatible quotient graphs where Dijkstra may be exact. |

**Table 3.**
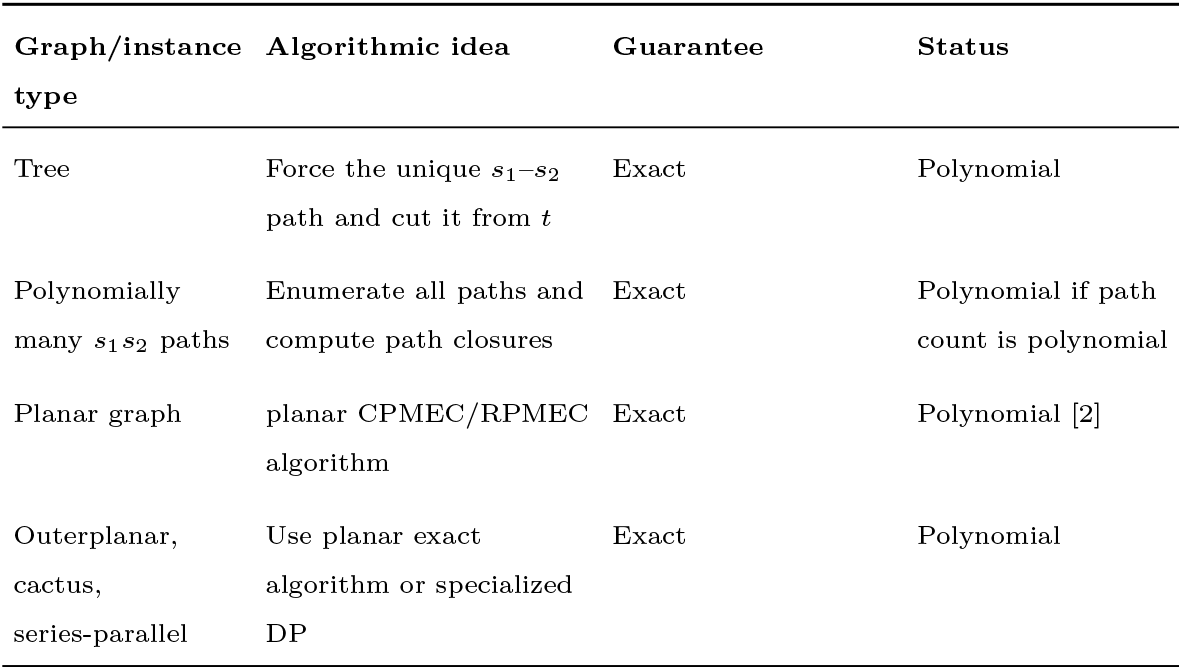
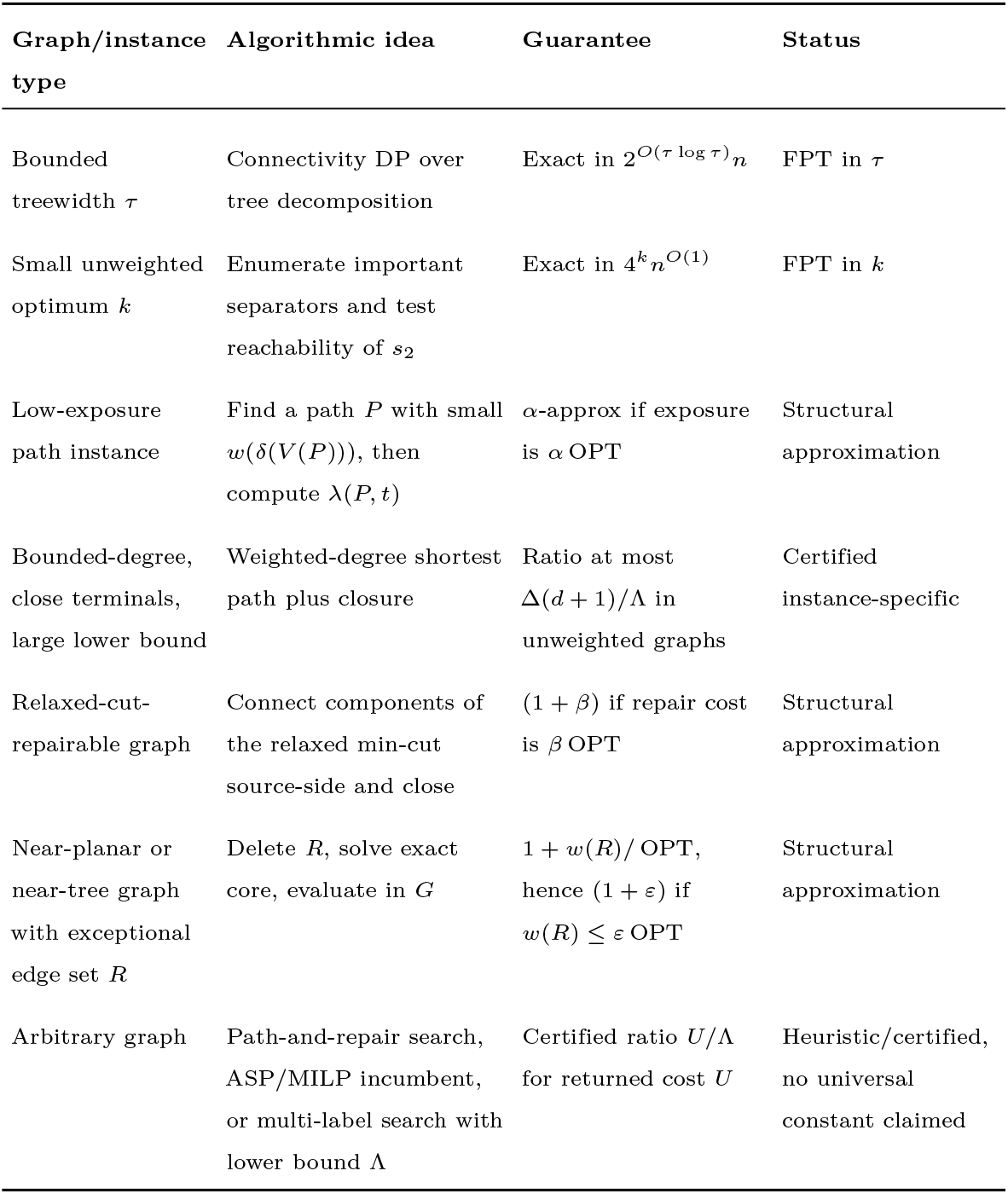
Approximation and exact algorithms for special classes of the three-terminal undirected edge RPMEC problem.

| Graph/instance type | Algorithmic idea | Guarantee | Status |
| --- | --- | --- | --- |
| Tree | Force the unique $s_1$–$s_2$ path and cut it from $t$ | Exact | Polynomial |
| Polynomially many $s_1 s_2$ paths | Enumerate all paths and compute path closures | Exact | Polynomial if path count is polynomial |
| Planar graph | planar CPMEC/RPMEC algorithm | Exact | Polynomial [2] |
| Outerplanar, cactus, series-parallel | Use planar exact algorithm or specialized DP | Exact | Polynomial |
| Bounded treewidth $\tau$ | Connectivity DP over tree decomposition | Exact in $2^{O(\tau \log \tau)} n$ | FPT in $\tau$ |
| Small unweighted optimum $k$ | Enumerate important separators and test reachability of $s_2$ | Exact in $4^k n^{O(1)}$ | FPT in $k$ |
| Low-exposure path instance | Find a path $P$ with small $w(\delta(V(P)))$, then compute $\lambda(P, t)$ | α-approx if exposure is $\alpha$ OPT | Structural approximation |
| Bounded-degree, close terminals, large lower bound | Weighted-degree shortest path plus closure | Ratio at most $\Delta(d+1)/\Lambda$ in unweighted graphs | Certified instance-specific |
| Relaxed-cut-repairable graph | Connect components of the relaxed min-cut source-side and close | $(1+\beta)$ if repair cost is $\beta$ OPT | Structural approximation |
| Near-planar or near-tree graph with exceptional edge set $R$ | Delete $R$, solve exact core, evaluate in $G$ | $1 + w(R)/\text{OPT}$, hence $(1+\varepsilon)$ if $w(R) \leq \varepsilon \text{OPT}$ | Structural approximation |
| Arbitrary graph | Path-and-repair search, ASP/MILP incumbent, or multi-label search with lower bound $\Lambda$ | Certified ratio $U/\Lambda$ for returned cost $U$ | Heuristic/certified, no universal constant claimed |

### Biological Pathway Graphs Where the Dijkstra-Style Algorithm Is Exact

The RPMEC formulation is natural in computational biology: one may want to block reachability to an undesirable state *t*, such as a disease-associated module, toxic byproduct, viral process, or adverse phenotype, while preserving reachability between protected biological functions *s*_1_ and *s*_2_. For example, *s*_1_ may represent an upstream regulatory signal and *s*_2_ an essential downstream response that should remain functional after an intervention. The key point of the preceding sections is that the Dijkstra-style algorithm is not justified merely because a biological graph is sparse, acyclic, or visually pathway-like. It is justified when the source-side optimum regions are frontier-realizable.

Biological pathway data are heterogeneous. KEGG pathway maps are manually drawn molecular interaction, reaction, and relation diagrams [18]. Reactome contains both a hierarchical event organization and detailed reaction diagrams; Reactome reaction diagrams need not be linear and may be branched or circular [19, 32]. Protein-protein interaction networks may have small-world and hub-rich structure [33, 34]. Metabolic reaction graphs can be compressed by contracting strongly connected components into metabolic building blocks, producing a metabolic DAG [22]. Consequently, the correct practical statement is not that all biological pathway graphs are easy, but that several biologically meaningful representations produce planar, laminar, arborescent, or dominator-compatible instances.

#### Module-respecting biological RPMEC instances

Many biological interventions are naturally module-respecting: one usually does not want to cut inside a stable protein complex, an inseparable feedback loop, a reversible metabolic block, or a curated subpathway. Such modules can be contracted before solving RPMEC.

Let 𝒫 = *{M*_1_, …, *M*_*r*_*}* be a partition of the biological graph into modules. A cut is *module-respecting* if it cuts only inter-module edges, or equivalently if every module lies completely on one side of the source-side cut. The quotient graph *H* = *G/*𝒫 has one vertex for each module and an edge or arc whenever the original graph has an interaction between the corresponding modules. The terminals *s*_1_, *s*_2_, *t* are mapped to their modules *M* (*s*_1_), *M* (*s*_2_), *M* (*t*).

##### Theorem 21

(Biological module-quotient sufficient condition). *Suppose the biological RPMEC instance is restricted to module-respecting cuts. If the quotient graph H* = *G/*𝒫 *satisfies the undirected or directed frontier-realizability condition of the previous sections, then the Dijkstra-style algorithm returns an optimal module-respecting RPMEC cut for the original graph. In particular, this holds when H is an undirected planar graph, an undirected tree/cactus/outerplanar graph, a directed out-arborescence rooted at M* (*s*_1_), *or a directed graph whose canonical optimal source-side regions are laminar and dominator-compatible*.

*Proof*. Every module-respecting source-side set *X* ⊆ *V* (*G*) corresponds exactly to a source-side set 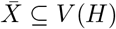 (*H*) in the quotient: a module is in 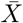 if and only if all of its vertices are in *X*. The inter-module cut cost in *G* is the same as the cut cost of 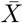 in *H*, after assigning each quotient edge the total cost of the original edges between the two modules. Reachability between *s*_1_ and *s*_2_ is also preserved at the quotient level, because internal module connectivity is treated as preserved or assigned infinite cost. Therefore the module-respecting RPMEC problem on *G* is equivalent to RPMEC on *H*. If *H* satisfies frontier-realizability, the Dijkstra-style algorithm is exact on *H* by the previous characterization. Lifting the quotient cut back to its original inter-module edges gives an optimal module-respecting cut in *G*.

#### Example class 1: undirected planar curated pathway maps

A curated pathway map can be converted into an undirected interaction graph by treating genes, proteins, complexes, reactions, or metabolites as vertices and treating physical interactions, regulatory relations, or reaction adjacencies as edges. Suppose the extracted graph is planar after biologically justified preprocessing, such as removing ubiquitous currency-metabolite shortcuts or contracting stable complexes. Then the undirected Dijkstra-style RPMEC algorithm is exact by the planar CPMEC result.

A representative use case is the following. Let *s*_1_ be an upstream receptor or regulatory protein, let *s*_2_ be an essential downstream transcription factor or metabolic product, and let *t* be a disease-associated output or toxic branch. The goal is to remove a minimum-cost set of pathway interactions that separates the source-side biological function from *t* while keeping the desired signal from *s*_1_ to *s*_2_. If the resulting graph is planar, every optimal protected region is forced to expand through a noncrossing frontier, so the Dijkstra-style relaxation cannot miss a cheaper hidden optimum.

The important qualification is that the map must be planar as a graph, not merely drawn without obvious crossings in a database image. This can be certified by running a standard planarity test on the extracted graph. If the graph is nonplanar because of a small number of cross-talk edges, one may still obtain an exact Dijkstra-solvable instance by contracting or deleting biologically irrelevant edges only when this is justified by the modeling assumptions.

#### Example class 2: Reactome-style pathway hierarchy trees

Reactome organizes biological events in a nested hierarchy of pathways, subpathways, and reactions [19, 32]. If RPMEC is posed on this hierarchy rather than on the full molecular reaction diagram, the graph is often a rooted tree or a nearly tree-like hierarchy. In such a model, a source-side feasible region is a rooted connected subtree, and the canonical optimum regions are nested. Therefore the laminar sufficient condition applies.

For example, *s*_1_ may be a high-level pathway event representing activation of a protective immune or stress-response program, *s*_2_ may be a required downstream subpathway, and *t* may be a subpathway associated with an adverse response. The Dijkstra-style algorithm is exact when the hierarchy has a single parent-child route from *s*_1_ to each relevant subpathway. The computed cut then identifies the minimum-cost parent-child links that block the adverse branch while preserving the desired branch.

The caveat is that Reactome’s detailed reaction diagrams are not necessarily trees: they may be branched or circular, and reactions may have multiple preceding or subsequent reactions [19]. Thus the guarantee applies automatically to the hierarchy abstraction, but the detailed reaction-level graph must be tested separately for planarity, laminarity, or dominator compatibility.

#### Example class 3: directed signaling cascades with dominator structure

A directed signaling pathway is often abstracted as a cascade

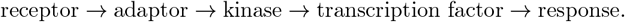

If the relevant directed graph is an out-arborescence rooted at *s*_1_, or more generally if each important vertex has a unique dominator chain from *s*_1_, then the directed Dijkstra-style algorithm is exact. Every feasible region that reaches a downstream vertex must contain the same upstream prefix, and hence the next optimum region is exposed by a single outgoing frontier arc.

A typical instance is: *s*_1_ is an activated receptor or upstream kinase, *s*_2_ is a beneficial transcriptional response that must remain reachable, and *t* is an off-target inflammatory, apoptotic, or oncogenic output. Edge weights may encode intervention cost, confidence penalty, or expected toxicity of blocking a relation. If cross-talk has been removed or modeled as high-cost non-cuttable structure, the quotient graph behaves as a cascade or arborescence, and the Dijkstra-style algorithm returns the optimum intervention set.

This class should not be confused with arbitrary directed pathway DAGs. The preceding NP-completeness result shows that a directed acyclic pathway can still contain set-cover-like global choices. The safe structure is not acyclicity alone, but single-frontier reachability induced by an arborescence or dominator-compatible DAG.

#### Example class 4: SCC-contracted feedback modules

Biological pathways often contain feedback loops. In a directed graph, a feedback loop appears as a strongly connected component. If the intervention should preserve the loop as an intact regulatory module, then the SCC can be contracted to a supernode. The Dijkstra-style algorithm can then be applied to the quotient when the quotient is a tree, outerplanar graph, planar graph, or dominator-compatible directed graph.

For example, a feedback-regulated kinase module may have several internal phosphorylation and dephosphorylation relations. Cutting inside that module may be biologically undesirable or experimentally infeasible. The module can be assigned infinite internal cut cost and contracted. RPMEC then chooses inter-module cuts that separate the bad-state module *t* while preserving reachability from the input module *s*_1_ to the essential module *s*_2_. Exactness follows from Theorem 21 once the quotient passes one of the structural tests.

#### Example class 5: certified metabolic m-DAGs

Metabolic networks are often represented through reaction graphs. A useful preprocessing step is to contract strongly connected reaction blocks into metabolic building blocks (MBBs). The condensation of the reaction graph is a metabolic DAG, or m-DAG [22]. This representation can reveal linear metabolic paths and cut nodes in the pathway, while keeping information about connectivity and robust reaction blocks.

An RPMEC instance on an m-DAG may take *s*_1_ to be a nutrient-entry reaction block, *s*_2_ an essential product or biomass-precursor block, and *t* a toxic byproduct branch or undesired disease-associated metabolic state. The Dijkstra-style algorithm is exact if the m-DAG is not just acyclic but also frontier-realizable, for example if it is an out-arborescence, a single-entry/single-exit module DAG, or a dominator-laminar DAG. In that case, preserving reachability from *s*_1_ to *s*_2_ forces a unique or nested sequence of metabolic building blocks, and the optimum cut cannot be hidden behind incompatible fronts.

The caveat is crucial: arbitrary m-DAGs are still DAGs, and DAG-RPMEC remains NP-complete. Thus an m-DAG should be certified by dominator or laminarity tests before the Dijkstra-style algorithm is claimed to be exact. If the m-DAG has extensive branching with reconvergence, an exact ASP or integer-programming formulation is safer.

#### Example class 6: functional-module quotients of protein-protein interaction networks

Raw protein-protein interaction networks are usually not safe instances for the Dijkstra-style algorithm. They may have small-world connectivity and hub-rich degree distributions [33, 34]. Hubs and shortcuts can create overlapping source-side regions, exactly the kind of structure that violates frontier-realizability.

However, PPI-derived RPMEC instances can become Dijkstra-solvable after module extraction. Suppose proteins are clustered into functional modules such as a repair module, signaling module, transcriptional module, or disease module, and the quotient graph between modules is a tree, cactus, outerplanar graph, planar graph, or directed arborescence. Then the Dijkstra-style algorithm is exact on the module quotient. In a disease-module example, *s*_1_ may represent a normal upstream regulator, *s*_2_ a required housekeeping or repair module, and *t* an oncogenic or pathogenic module. The cut identifies low-cost inter-module interactions whose removal blocks the pathogenic module while preserving the protected functional route.

This distinction is important in applications. The claim is not that whole-cell PPI networks are easy. The claim is that biologically meaningful module-tree or module-planar quotients are exact Dijkstra-solvable RPMEC instances.

#### Comparison of biological graph classes

**Table 4.**
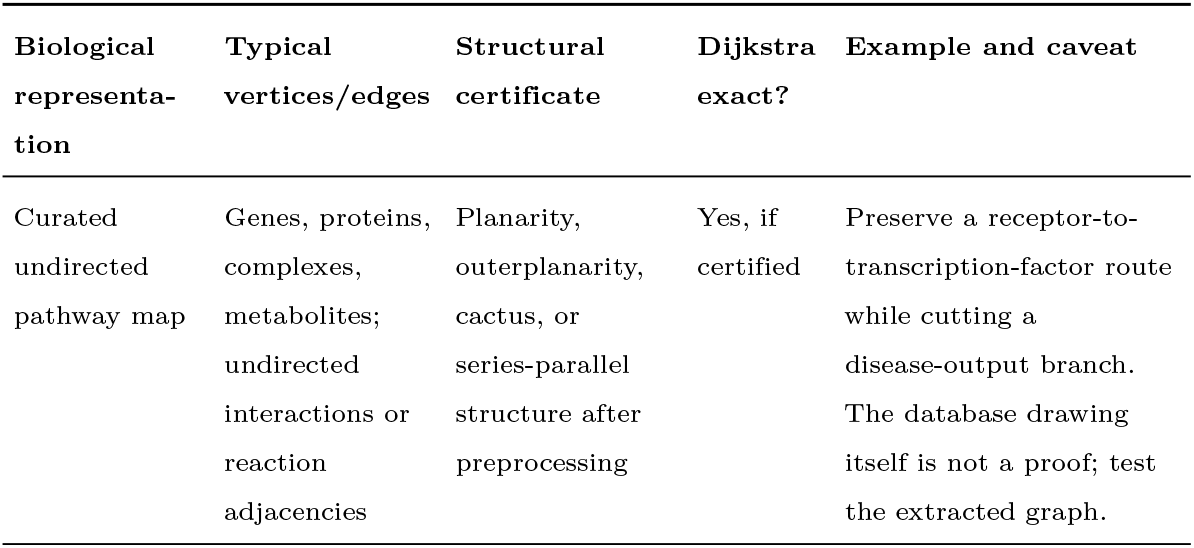
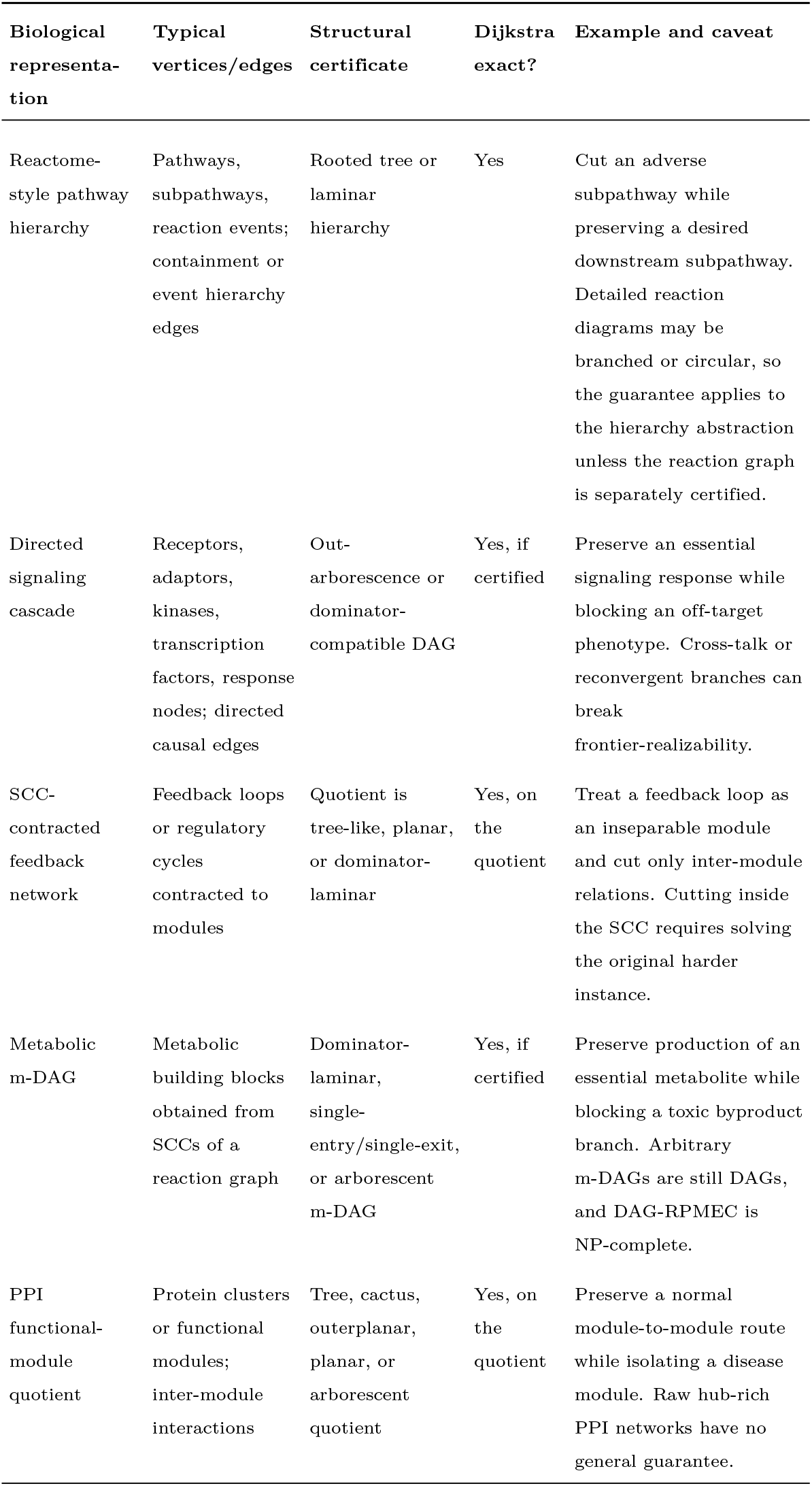
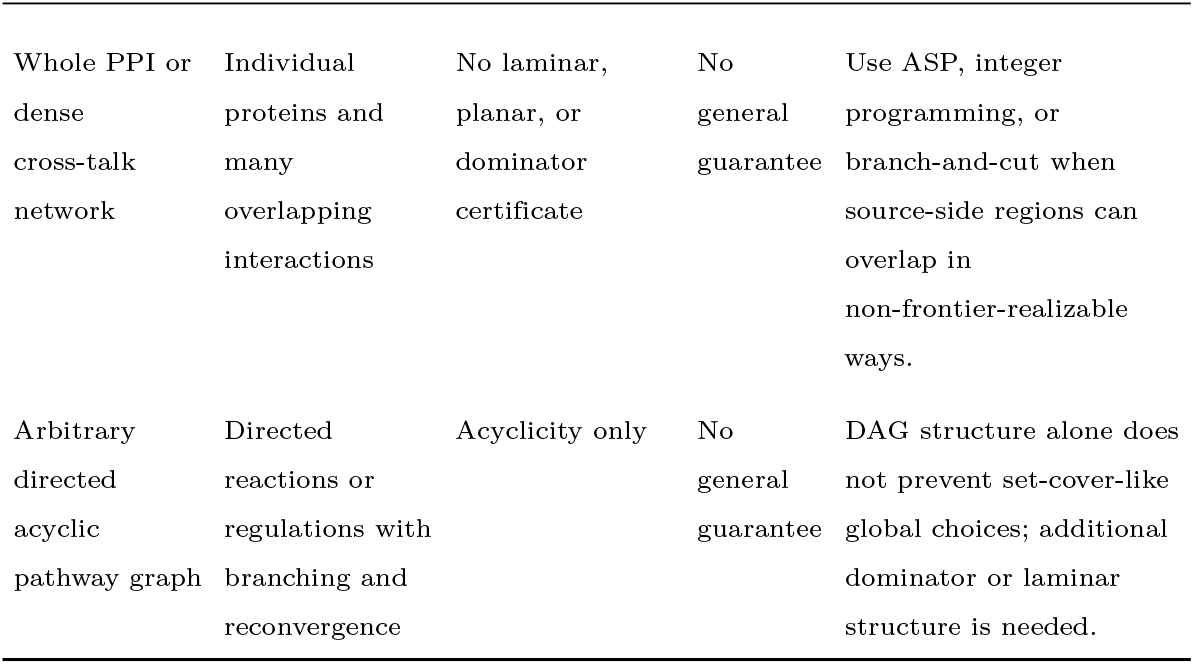
Practical biological graph classes for RPMEC. The Dijkstra-style algorithm is exact only for representations whose extracted or contracted graph satisfies a frontier-realizability certificate.

| Biological representation | Typical vertices/edges | Structural certificate | Dijkstra exact? | Example and caveat |
| --- | --- | --- | --- | --- |
| Curated undirected pathway map | Genes, proteins, complexes, metabolites; undirected interactions or reaction adjacencies | Planarity, outerplanarity, cactus, or series-parallel structure after preprocessing | Yes, if certified | Preserve a receptor-to-transcription-factor route while cutting a disease-output branch. The database drawing itself is not a proof; test the extracted graph. |
| Reactome-style pathway hierarchy | Pathways, subpathways, reaction events; containment or event hierarchy edges | Rooted tree or laminar hierarchy | Yes | Cut an adverse subpathway while preserving a desired downstream subpathway. Detailed reaction diagrams may be branched or circular, so the guarantee applies to the hierarchy abstraction unless the reaction graph is separately certified. |
| Directed signaling cascade | Receptors, adaptors, kinases, transcription factors, response nodes; directed causal edges | Out-arborescence or dominator-compatible DAG | Yes, if certified | Preserve an essential signaling response while blocking an off-target phenotype. Cross-talk or reconvergent branches can break frontier-realizability. |
| SCC-contracted feedback network | Feedback loops or regulatory cycles contracted to modules | Quotient is tree-like, planar, or dominator-laminar | Yes, on the quotient | Treat a feedback loop as an inseparable module and cut only inter-module relations. Cutting inside the SCC requires solving the original harder instance. |
| Metabolic m-DAG | Metabolic building blocks obtained from SCCs of a reaction graph | Dominator-laminar, single-entry/single-exit, or arborescent m-DAG | Yes, if certified | Preserve production of an essential metabolite while blocking a toxic byproduct branch. Arbitrary m-DAGs are still DAGs, and DAG-RPMEC is NP-complete. |
| PPI functional-module quotient | Protein clusters or functional modules; inter-module interactions | Tree, cactus, outerplanar, planar, or arborescent quotient | Yes, on the quotient | Preserve a normal module-to-module route while isolating a disease module. Raw hub-rich PPI networks have no general guarantee. |
| Whole PPI or dense cross-talk network | Individual proteins and many overlapping interactions | No laminar, planar, or dominator certificate | No general guarantee | Use ASP, integer programming, or branch-and-cut when source-side regions can overlap in non-frontier-realizable ways. |
| Arbitrary directed acyclic pathway graph | Directed reactions or regulations with branching and reconvergence | Acyclicity only | No general guarantee | DAG structure alone does not prevent set-cover-like global choices; additional dominator or laminar structure is needed. |

## Conclusion

We studied the three-terminal Reachability Preserving Minimum Edge Cut problem, where the goal is to separate two protected source nodes from a target node while keeping the protected nodes connected. The problem differs fundamentally from ordinary minimum cut because the feasible source-side must satisfy both a separation constraint and a connectivity-preservation constraint. We revisited the Dijkstra-style algorithm originally motivated by planar instances and showed that its success depends on a stronger structural property: every next optimal source-side region must be discoverable through a one-frontier relaxation. This frontier-realizability condition explains why the algorithm is exact on planar, laminar, tree-like, and other structurally accessible instances, while it may fail on general graphs.

We also considered broader algorithmic directions for the original three-terminal edge case. The open complexity of the general undirected edge version appears closely related to cut-uncut and network-diversion variants. For exact computation, we discussed ASP and MILP formulations, bounded-treewidth dynamic programming, important-separator-based fixed-parameter algorithms, and multi-label source-region search. For approximation, we developed a path-closure viewpoint in which each protected *s*_1_-to-*s*_2_ path induces a standard min-cut subproblem. This leads to certified approximation heuristics and stronger guarantees on low-exposure, near-planar, near-tree, and relaxed-cut-repairable graphs.

Finally, we identified biological pathway and module graphs as a practical domain where the structural theory is useful. Curated pathway maps, hierarchy trees, SCC-contracted feedback modules, metabolic building-block DAGs, and functional-module quotient graphs may yield planar, laminar, or dominator-compatible structure. On such instances, the Dijkstra-style algorithm can be used as an exact solver after suitable certification or preprocessing. On dense cross-talking networks, however, exact ASP, MILP, or parameterized methods remain more appropriate. Future work includes proving or refuting NP-hardness of the original undirected three-terminal edge case, developing stronger approximation guarantees, and evaluating certified algorithms on real biological and cyber-security networks.

## Acknowledgments

Author information, funding, data availability, and author contributions will be completed during final submission preparation.

